# Transcriptome-wide profiling of alternative splicing regulators with CRISPore-seq

**DOI:** 10.1101/2025.11.25.690515

**Authors:** Simon Müller, Nathanael Andrews, Rachel E. Yan, Akash Sookdeo, Wells H. Burrell, Xiaoguang Dai, Priyesh Rughani, Zharko Daniloski, Sissel Juul, Neville E. Sanjana

## Abstract

Alternative splicing creates diverse RNA isoforms from individual genes, yet single-cell CRISPR screens are limited to gene-level quantification and cannot detect changes in alternative splicing and transcript isoforms. To overcome this limitation, we develop CRISPore-seq, which couples massively-parallel CRISPR perturbations with joint short- and long-read transcriptomics. CRISPore-seq simultaneously captures genetic perturbations and expression of genes, full-length transcripts and surface proteins in single cells. CRISPore-seq long reads identify 80% more transcript isoforms than short reads. Nearly all long reads map to unique transcript isoforms — in contrast to existing single-cell perturbation methods, which rarely distinguish specific isoforms. Using CRISPore-seq, we knock-down 15 different RNA-binding proteins (RBPs) and identify thousands of perturbation-driven alternative splicing events (ASEs). We find that exon skipping is the most common ASE observed and that skipped exons are enriched for binding sites of perturbed RBPs. Loss of the Nager syndrome-associated spliceosomal factor *SF3B4* triggers skipping of exon 2 in the cell-cycle regulator *CCND1*, preventing formation of a complex with CDK6 and blocking the G1-S transition. After rescue with a *CCND1* isoform containing the skipped exon, both holoenzyme complex formation and cell proliferation are restored. By linking genes to transcriptional phenotypes with isoform-level resolution, CRISPore-seq is a highly scalable tool for understanding the impact of genetic perturbations on the human transcriptome.

## Introduction

Alternative splicing (AS) of pre-messenger RNAs is a post-transcriptional mechanism to expand transcript diversity in mammalian cells^1–3^. During splicing, introns are removed from pre-mRNA and exons are joined together. However, AS allows for different combinations of exons to be included or excluded^4^. Introns or parts of them can also be retained. In human cells, over 90% of multi-exon genes undergo alternative splicing^5^. RNA-binding proteins (RBPs), which regulate alternative splicing, generate multiple distinct mRNA variants from a single gene. These isoforms can encode proteins with distinct cellular functions. Isoform diversity is fundamental for biological processes, such as development and tissue homeostasis^6–9^. Alterations in AS can cause or influence the progression of many human diseases, including cancer and neurodegenerative disorders^10,11^.

Recent large-scale efforts have mapped AS landscapes and identified regulatory roles of RBPs. These studies profiled transcriptomes after perturbing individual RBPs or core spliceosome components, providing comprehensive atlases^12,13^. However, arrayed perturbations and bulk RNA sequencing inherently limit their scalability; these approaches also cannot measure variability between individual gene-perturbed cells and have poor sensitivity to global transcriptional changes due to normalization between different perturbations.

Pooled CRISPR perturbation screens allow large-scale gene function studies across many cell types^14–17^. Combining these screens with single-cell RNA sequencing (scRNA-seq) enables high-throughput profiling of transcriptional responses to genetic perturbations in individual cells^18–21^. However, common short-read scRNA-seq methods usually only profile transcript ends (3′ or 5′). This limitation makes it difficult for computational methods to accurately reconstruct isoforms. Distinguishing between different isoforms, which can have distinct functions, remains challenging due to low read coverage and sparsity of data.

To overcome these limitations, we developed CRISPore-seq, which combines scalable pooled CRISPR perturbations with joint short-read and long-read transcriptomics. We use this method to perturb known alternative splicing regulators in human cells and quantify their effect on gene and isoform expression in single cells. We map RBP-specific transcriptional phenotypes and pinpoint affected cellular pathways. Furthermore, we leverage long-read transcriptomics to discover functional alternative splicing events.

## Results

### Single-cell CRISPR screens with long-read transcriptomics

We sought to study transcriptional phenotypes and isoform diversity in single cells, driven by perturbing various alternative splicing regulators. For this, we developed CRISPore-seq, a scalable platform that uses long-read nanopore sequencing to achieve full-length transcript coverage (**Fig. 1a**). It builds on Enhanced CRISPR-compatible Cellular Indexing of Transcriptomes and Epitopes with single-cell sequencing (ECCITE-seq)^22^, which combined pooled CRISPR screens with short-read scRNA-seq to profile gene expression, CRISPR perturbations and cell surface markers.

**Fig. 1:**
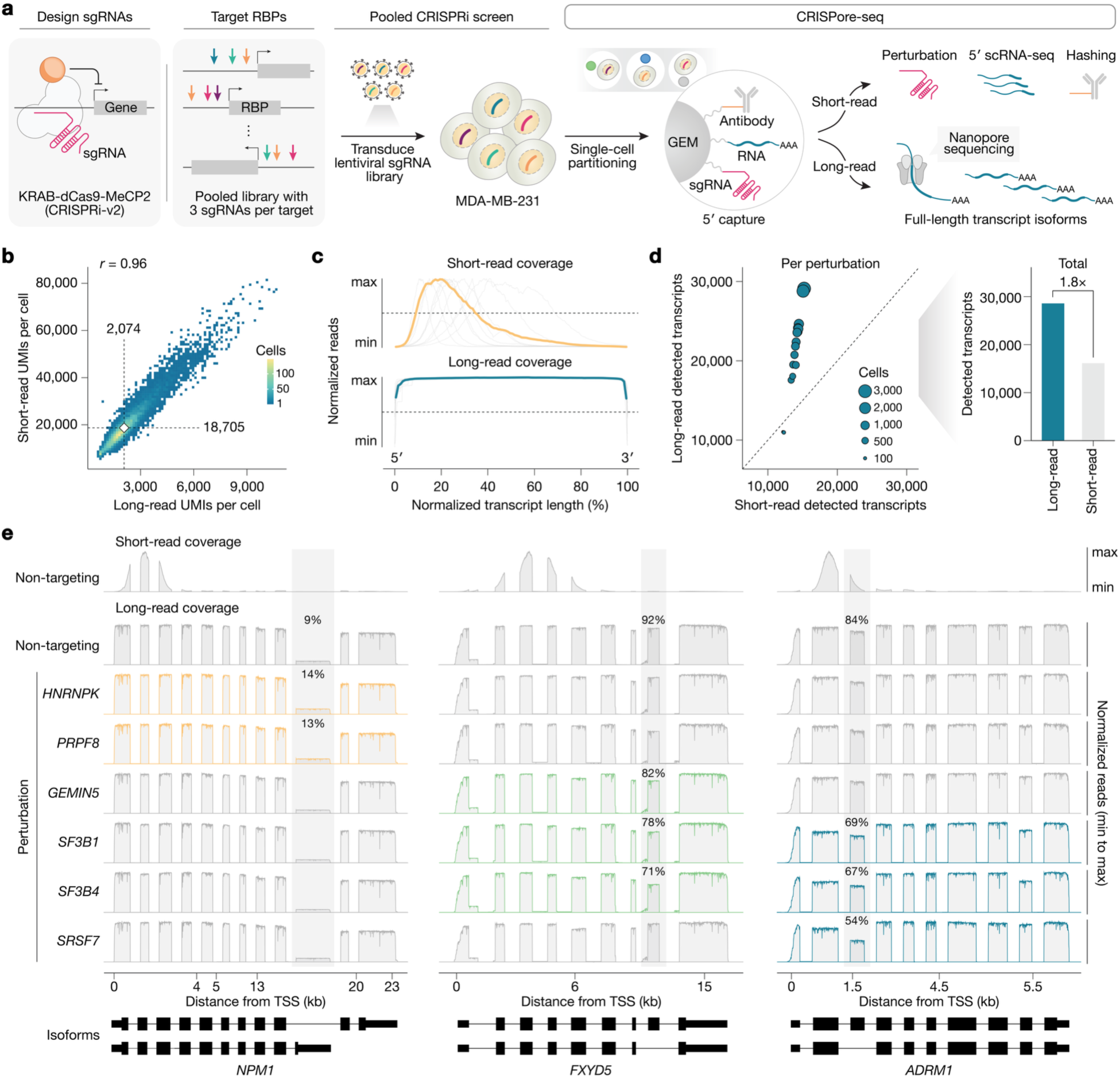
Pooled CRISPR knockdown coupled with single-cell short- and long-read transcriptomics. **a,** CRISPore-seq combines short and long-read sequencing to reveal isoform changes in full-length transcripts, while capturing sgRNA, RNA and oligo-conjugated antibodies. **b,** Pearson correlation between unique molecular identifiers (UMIs) per cell as measured by short- and long-read sequencing. **c,** Normalized read coverage from short- and long-read sequencing for ten highly-expressed transcripts (*gray*). Line fits estimated using LOESS regression (*yellow* and *blue*). **d,** Total detected protein-coding transcripts across perturbations with short- and long-read sequencing. **e,** Isoforms of *NPM1*, *FXYD5*, and *ADRM1* detected using short- and long-read sequencing in cells with perturbations targeting RNA-binding proteins *GEMIN5*, *HNRNPK*, *PRPF8*, *SF3B1*, *SF3B4*, *SRSF7*, and cells with non-targeting control perturbations. Reads are normalized to total reads at the locus and with introns compressed for visualization (see *Methods*).

First, we engineered a human breast cancer cell line to express a dual-effector CRISPR inhibition system, KRAB-dCas9-MeCP2 (CRISPRi-v2)^21^. Next, we created a single-guide RNA (sgRNA) library to perturb diverse RBPs with three sgRNAs targeting the transcription start sites of each RBP (**Supplementary Table 1**). We also added negative control (non-targeting) perturbations designed to not target anywhere in the human genome.

We transduced this pooled lentiviral sgRNA library at a low multiplicity-of-infection (∼0.05). This ensured that most cells received a single CRISPR perturbation. Next, we collected the perturbed cells. We used droplet-based microfluidics for single-cell partitioning and prepared CRISPore-seq libraries (**Fig. 1a**). Short-read sequencing identified captured sgRNAs (perturbations), RNA transcripts, and oligo-conjugated antibodies. Long-read nanopore sequencing characterized full-length transcript isoforms. Notably, oligo-conjugated antibodies enabled superloading and multiplet discrimination (cell hashing). However, they could also be used to profile the expression of cell surface protein markers^22^. In this manner, we achieve both high-depth coverage (short-read) and accurate isoform determination (long-read) in the same single cells.

Next, we assessed the performance and complementarity of short- and long-read sequencing by comparing their quality metrics. Cell barcode detection showed high concordance, with ∼92% of high-quality barcodes present in both datasets (**Supplementary Fig. 1a**). The number of unique molecular identifiers (UMIs) per cell and per gene between the two sequencing modalities were well correlated (*r* = 0.96) (**Fig. 1b and Supplementary Fig. 1b**). We then analyzed differential gene expression in both short-read and long-read data by fitting read counts to a negative binomial distribution and using permutation tests optimized for single-cell perturbation data^23^. For differentially expressed genes (DEGs) detected in either modality, transcript changes also showed a high correlation (*r* = 0.86) with minimal discordance (<3%) (**Supplementary Fig. 1c**).

We then assessed isoform resolution, the key advantage of long-read sequencing over conventional short-read methods. As expected, short-read sequencing reads only resolved the 5′ end of highly expressed transcripts (**Fig. 1c**). By their midpoint, less than 50% of the transcripts had at least 5% read coverage. In contrast, long-read sequencing achieved complete 5′-to-3′ coverage and identified 80% more unique protein-coding transcript isoforms across all perturbations (**Fig. 1d**). Overall, long-read transcript sequencing resolved 85% of isoforms, whereas short-read sequencing reads could distinguish only 20% (**Supplementary Fig. 1d**).

For instance, long-read sequencing detected two major isoforms of *NPM1*. These differed at their 3′ ends due to alternative termination sites and 3′ untranslated regions (**Fig. 1e**). Short-read sequencing, primarily resolving the 5′ end, could not distinguish these isoforms. Therefore, only long-read sequencing could detect *NPM1* isoform changes after RBP perturbations. Following *HNRNPK* and *PRPF8* knockdowns, the shorter *NPM1* isoform increased 1.5- and 1.4-fold, respectively, compared to cells with non-targeting perturbations. Perturbations of *SF3B1/4*, *GEMIN5*, and *SRSF7* showed no difference in *NPM1* isoform expression. We also observed distinct RBP-driven changes in other isoforms. This included exon skipping in late exons of *FXYD5* and early exons of *ADRM1* (**Fig. 1e**). Thus, CRISPore-seq not only enables the profiling of CRISPR perturbations and overall gene expression but also resolves full-length transcript changes linked to genetic perturbations.

### Transcriptional phenotypes of RNA-binding proteins

RBPs play diverse roles in post-transcriptional gene regulation^24,25^. To explore their specific functions, we used CRISPore-seq to interrogate RBPs with varying essentiality, molecular functions, and domain architectures (**Fig. 2a**)^12,24,26^. We identified distinct expression patterns triggered by RBP perturbations in single cells, while confirming on-target knockdown across perturbations (**Fig. 2b and Supplementary Fig. 1e**). As a control, we compared cells receiving different non-targeting sgRNAs and found no differentially expressed genes (**Supplementary Fig. 1f**).

**Fig 2:**
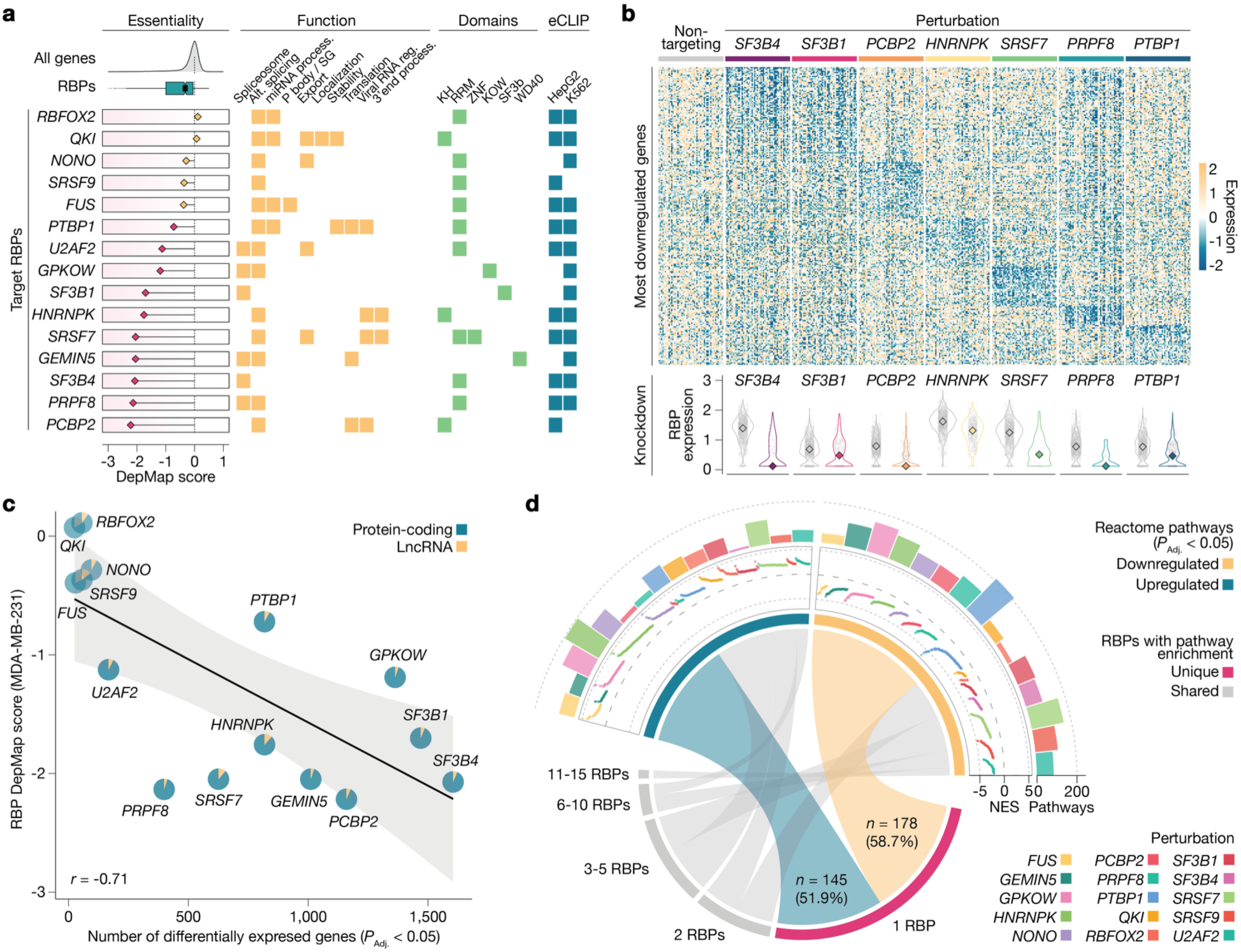
Transcriptional phenotypes of RNA-binding protein perturbations. **a,** RNA-binding proteins targeted using CRISPore-seq by their essentiality, function, domains, and enhanced Crosslinking and Immunoprecipitation (eCLIP) profiling. **b,** Single-cell expression heatmap from short-read sequencing showing the 25 most downregulated genes for 50 randomized cells per selected target perturbation (*top*) and the target gene expression (*bottom*). **c,** Pearson correlation between DepMap essentiality score and number of differentially expressed genes (lncRNAs and protein-coding genes, *P*_adj._ < 0.05) following RBP perturbations from short-read sequencing. Pie charts show the fraction of differentially-expressed lncRNAs and protein-coding genes. **d,** Altered Reactome pathways in cells with RBP perturbations. Outer tracks show the number of significant pathways per RBP (*P*_adj._ < 0.05) and normalized enrichment scores (NES). The central chord diagram shows pathway overlaps among RBP perturbations. Pathway significance was assessed by a Kolmogorov-Smirnov test (Benjamini-Hochberg corrected).

We mapped single-cell transcriptomes to a two-dimensional embedding using short-read sequencing (**Supplementary Fig. 1g**). Cells lacking the essential RBP *SRSF7*, an SR-rich splicing factor, formed a distinct cluster. *SRSF7* knockdown broadly altered gene expression, including ∼600 DEGs and downregulation of core spliceosome components (**Supplementary Fig. 1h, i**). Conversely, knocking down the non-essential RBP *NONO* had only a modest effect on gene expression and the splicing machinery. We then analyzed transcriptional phenotypes across all RBP perturbations. RBP essentiality significantly correlated with the number of RBP-driven DEGs (*r* = -0.71) (**Fig. 2c, Supplementary Fig. 1j and Supplementary Table 2**). Thus, loss of highly essential RBPs triggers broad changes in gene expression, primarily affecting protein-coding genes that then alter cellular fitness phenotypes.

Next, we sought to understand how different RBP perturbations affect gene programs. To annotate gene functions, we performed gene set enrichment analysis (GSEA) using two curated pathway annotations from the Gene Ontology (GO) and Reactome databases^27,28^. Across all RBP perturbations, 888 GO and 582 Reactome pathways were significantly affected (*P*_Adj._ < 0.05, **Fig. 2d**, **Supplementary Fig. 2a and Supplementary Table 3**). Notably, across both the Reactome and GO analyses, most significant pathways tended to be specific to individual RBPs (55% of Reactome pathways, 74% of GO pathways) and not widely shared (**Fig. 2d and Supplementary Fig. 2b**), although we observed higher correlation after knockdown of essential RBPs (those causing fitness defects), and similarly, after non-essential RBP knockdowns (**Supplementary Fig. 2c**). Furthermore, GSEA from short- and long-read transcriptomics were highly correlated (**Supplementary Fig. 3d**), highlighting the complementarity of sequencing approaches.

Many perturbations led to alterations in cell cycle-related pathways (**Supplementary Table 3**). Thus, we used the transcriptome data (from short- and long-read sequencing) to assign cell cycle phases for individual perturbed cells (**Supplementary Fig. 2e**). Various RBP perturbations impaired cell cycle progression, causing cells to accumulate in specific phases. For example, *PTBP1*-perturbed cells accumulated in G2M, with an impaired G1 transition. *RBFOX2*-perturbed cells stalled in S phase. We also found a frequent accumulation of cells in G1, particularly after knockdown of essential RBPs like *SF3B1*, *SRSF7*, *PRPF8* and *SF3B4*. For instance, after *SF3B4* knockdown, the fraction of cells in G1 phase increased ∼2-fold compared to cells that received non-targeting controls. Overall, the cell cycle alterations were consistent across both short- and long-read sequencing approaches (median *r* = 0.96, **Supplementary Fig. 2e**). Together, these findings show that CRISPore-seq effectively monitors transcriptome-wide expression changes and identifies associated cellular pathways following genetic perturbations, using both short- and long-read scRNA-seq.

### Decoding alternative splicing dynamics

To understand the transcript-level mechanisms driving these broader cellular phenotypes, we next focused on alternative splicing using long-read transcriptomics. We analyzed different AS events, including exon skipping, intron retention, and alternative splice site usage (3′ or 5′) (**Fig. 3a, Supplementary Fig. 3a and Supplementary Table 4**). Across all perturbations, we identified over 6,000 individual alternative splicing events. First, we assessed the magnitude of these changes. Perturbing core U2 snRNP components (*SF3B1* and *SF3B4*) caused global splicing changes. However, these changes were relatively modest across transcripts. In contrast, *SRSF7* and *PRPF8* perturbations resulted in fewer splicing changes, but these changes had a larger magnitude (**Fig. 3b and Supplementary Fig. 3a**).

**Fig. 3:**
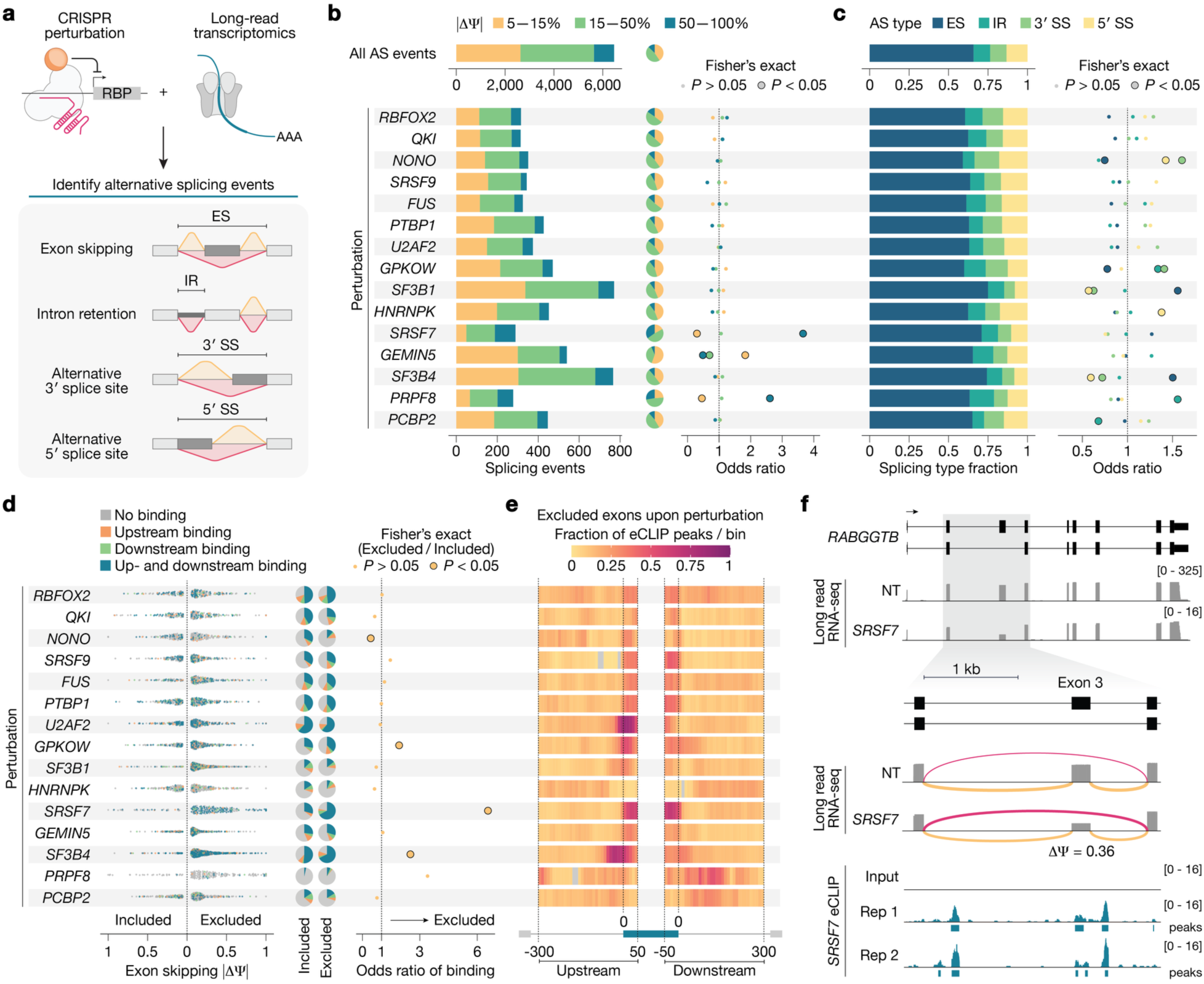
Alternative splicing patterns upon perturbation of RNA-binding proteins. **a,** CRISPore-seq combines CRISPR perturbations with long-read transcriptomics to identify alternative splicing (AS) events such as exon skipping (ES), intron retention (IR) and alternative 3′/5′ splice sites (SS). **b,** Number of alternative splicing (AS) events following RBP perturbations. AS events were grouped based on the percent spliced-in (ΔΨ). Cells were pseudo-bulked by perturbation and significant splicing events (|ΔΨ| > 0.05 and *P* < 0.05) were identified using FLAIR (see *Methods*). Bar plots show the number of total AS events captured per perturbation (*left*), pie charts show the relative fraction of each event (*middle*). The odds ratio and significance were computed by comparing the respective fraction per perturbation to the fraction across all AS events using Fisher’s exact test (*right*). **c,** Proportion of different AS events upon RBP perturbations (*left*). Enrichment and significance were calculated using Fisher’s exact test (*right*). **d,** Magnitude (|ΔΨ|) of exon skipping events, including exons that are included or excluded, after RBP knockdowns (*left*). Pie charts (*middle*) show RBP binding distribution in defined windows flanking ES exons (upstream: -300/+50 nt, downstream: -50/+300 nt) for ES events. Enrichment of RBP binding (odds ratio) and significance comparing included versus excluded exons (Fisher’s exact test) (*right*). **e,** Fraction of RBP binding within 10 nt windows in excluded exons upon RBP perturbations in upstream and downstream windows. **f,** *RABGGTB* splicing changes after *SRSF7* knockdown from long-read sequencing and eCLIP data showing *SRSF7* binding sites. Loops represent the relative frequency of exon inclusion (*yellow*) or exclusion (*pink*).

Overall, we found that exon skipping was the most frequent splicing alteration, making up 42-76% of changes depending on the perturbation (**Fig. 3c, Supplementary Fig. 3a**). For example, in cells with *SF3B1* and *SF3B4* perturbations, we found that the AS events were significantly enriched for exon skipping. In contrast, alternative splice site usage was depleted compared to the entire data set. Conversely, the perturbation of *NONO* led to more usage of alternative splice sites and fewer exon skipping events. Interestingly, *PRFP8* perturbation resulted in enriched intron retention events. *PCBP2* knockdowns, however, led to depleted intron retention. This highlights a key advantage of CRISPore-seq, which connects each RBP with its functional role in different transcripts.

Next, we combined our RBP-mediated alternative splicing data with ENCODE eCLIP binding^12,29^ (**Supplementary Table 5**). This allowed us to assess the correspondence between RBP binding and alternative splicing of the same transcript. Across all perturbed RBPs, AS events were enriched for transcripts with corresponding eCLIP peaks compared to random transcript sets (**Supplementary Fig. 3b**). First, we focused on exon skipping events and detected more excluded than included exons after RBP perturbations (**Fig. 3d and Supplementary Fig. 3c**). Next, we examined RBP binding in two distinct windows: upstream and downstream of alternatively spliced exons (**Supplementary Table 6**).

We then evaluated how RBPs influence their target RNA. For example, *SF3B1* and *SF3B4* perturbations both caused widespread exon exclusion. However, affected exons were more likely to overlap eCLIP peaks for *SF3B4* than for *SF3B1*, suggesting that *SF3B4* mediates more changes through direct binding (**Fig. 3d**). We also found that *SRSF7* binding is enriched around exons excluded upon its perturbation, indicating that *SRSF7* primarily promotes exon inclusion. Conversely, *NONO* binding indicated more exon inclusion after *NONO* knockdown.

Next, we focused on all excluded exons after RBP perturbation that showed direct RBP binding. We increased the resolution to 10 bp bins and determined the fraction with observed binding for each RBP (**Fig. 3e**). This analysis revealed distinct binding patterns: *SF3B4* showed dominant enrichment in the first 50 nucleotides (nt) of the upstream intron, while *U2AF2* primarily associated with the first 50 nt of the alternatively spliced exon. Other RBPs showed more diverse binding. *RBFOX2*, for example, tended to bind more frequently within the exon but also across the entire intronic range. *PCBP2* binding was more pronounced in the downstream intron of exons excluded after its perturbation. In contrast, *SRSF7* binding was dominant within the alternatively spliced exon (**Fig. 3e**). For example, *SRSF7* knockdown results in exon 3 skipping in *RABGGTB* (**Fig. 3f**). We found multiple *SRSF7* binding sites in the alternatively spliced exon and downstream intron, suggesting that *SRSF7* directly promotes the inclusion of this exon (**Fig. 3f**). Similarly, loss of *PTBP1* directly induced exon 2 skipping in *CFLAR* (**Supplementary Fig. 3d**). Together, these distinct RBP binding patterns around excluded exons indicate diverse, position-dependent mechanisms for regulating exon skipping.

We identified similar trends for intron retention events. Generally, more introns were retained after RBP perturbations compared to cells with non-targeting sgRNAs (**Supplementary Fig. 3e**). Many RBPs showed direct binding to these retained introns (**Supplementary Fig. 3e and Supplementary Table 7**). For instance, intron 2 of *RPS28* is retained after *SF3B1* and *SF3B4* perturbations, with direct *SF3B4* binding within the intron itself (**Supplementary Fig. 3f**), suggesting that *SF3B4* binding is necessary for the intron removal. Overall, RBP perturbations resulted in diverse alternative splicing events. The resulting transcript isoforms might, in turn, alter protein function.

### Exon skipping leads to loss-of-function isoforms

To study the functional impact of exon skipping, we combined RBP-driven alternative splicing data with exon-specific RNA-targeting CRISPR-Cas13 screens^30^ (**Fig. 4a**). We hypothesized that skipping specific exons after RBP perturbation would create loss-of-function isoforms. Thus, perturbing the exon-containing isoforms with Cas13 gRNAs might be critical for cell fitness. We found that perturbations of several transcripts led to marked depletion in pooled Cas13 screens compared to non-targeting guide RNAs (**Fig. 4b, Supplementary Fig. 4a and Supplementary Table 8**). For instance, targeting exon 3 of *RABGGTB* or exon 2 of *CCND1* with Cas13 caused depletion in the pooled screen. This exon of *CCND1* is skipped after *SF3B4* knockdown (change in percent spliced-in [ΔΨ] = 15%, **Fig. 4c**). This suggests the exon skipping event may contribute to the fitness phenotype observed with *SF3B4* perturbation. We also found that *SF3B4* directly binds to the introns upstream and downstream of this *CCND1* exon (**Fig. 4c**), indicating a direct effect on this alternative splicing event. Notably, while *SF3B4* knockdown specifically affected *CCND1* splicing, it did not change the overall gene expression of *CCND1* (log_2_ fold change = 0.06, **Supplementary Fig. 4b**). This highlights an important advantage of CRISPore-seq: Long-read sequencing resolves changes at isoform resolution that would be missed in conventional short-read scRNA-seq primarily reflecting gene-level abundance.

**Fig. 4:**
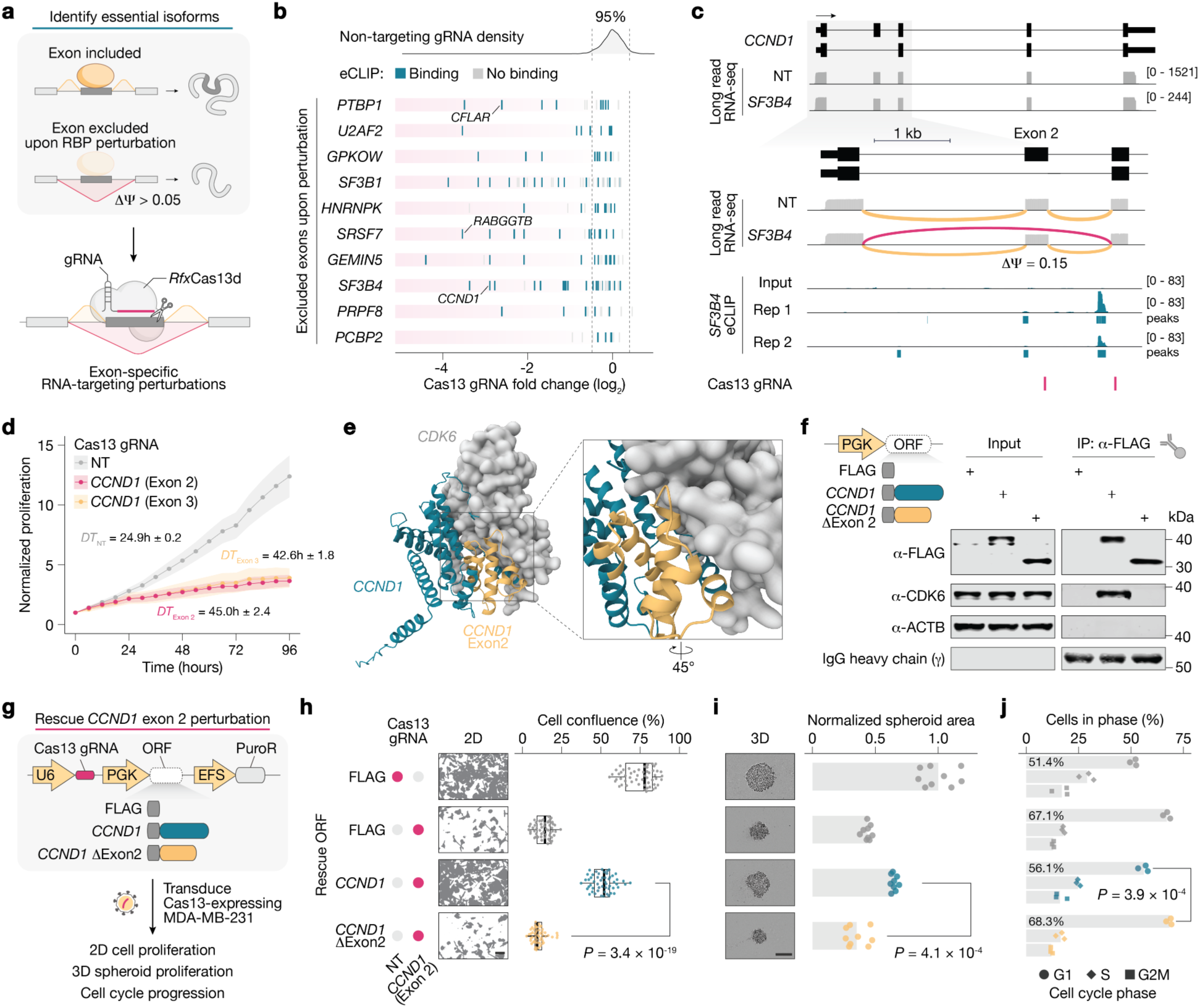
Exon skipping in *CCND1* leads to impaired cyclin-dependent kinase complex formation, impaired cell cycle progression and decreased proliferation. **a,** Cas13 targeting of isoform-specific exons to identify full-length isoforms critical for cell fitness. **b,** Fold change (day 14 vs. day 0) of individual gRNAs targeting the indicated genes with the 95% confidence interval of non-targeting gRNAs indicated by the *gray* lines. Cas13 data was gathered from gRNAs exclusively targeting excluded exons upon RBP perturbations in MDA-MB-231 cells^43^. **c,** *SF3B4* knockdown leads to *CCND1* exon 2 exclusion. Tracks display read coverage from CRISPore-seq (*above*) and eCLIP binding for *SF3B4* (*below*). Loops represent the relative frequency of exon inclusion (*yellow*) or exclusion (*pink*). The position of exon-specific Cas13 gRNAs is indicated in pink (*bottom*). **d,** Normalized proliferation of MDA-MB-231 cells transduced with Cas13 gRNAs targeting *CCND1* exon 2, or exon 3 and a non-targeting (NT) gRNA (*n* = 54 images per timepoint with three images per technical replicate, six technical replicates per transduction, and three independent transductions per perturbation). Points denote the mean and the shaded regions display the standard deviation. Doubling times (DT) are indicated as mean ± standard deviation. **e,** Structural predictions for the CCND1-CDK6 complex formation using AlphaFold3 ^31^. CCND1 exon 2-encoded amino acids (*yellow*) are highlighted at the interface with CDK6 (*gray*). **f,** Co-immunoprecipitation of CCND1 and CDK6 in HEK293FT cells transfected with expression vectors containing FLAG, FLAG-*CCND1* and FLAG-*CCND1*(ΔExon 2). Representative western blot images from 3 biological replicates. **g,** Overview of rescue experiment with Cas13-targeting of endogenous *CCND1* and overexpression of FLAG, FLAG-*CCND1*, and FLAG-*CCND1*(ΔExon 2). **h,** Proliferation (2D cell culture) of MDA-MB-231 cells after perturbation of endogenous *CCND1* and/or rescue with FLAG, FLAG-*CCND1*, or FLAG-*CCND1*(ΔExon 2) expression. Representative images show cells after 4 days of growth (*n* = 54 images with three images per technical replicate, three technical replicates per transduction, and three independent transductions per perturbation). Boxplots show median, interquartile range, and whiskers indicating 1.5× interquartile range. Significance was determined using a two-sided Mann-Whitney *U*-test. Scale bar, 100 μm. **i,** Spheroid proliferation assay (3D cell culture) of MDA-MB-231 cells after perturbation of endogenous *CCND1* and/or rescue with FLAG, FLAG-*CCND1*, or FLAG-*CCND1*(ΔExon 2) expression. Representative images show cells after 7 days of growth post induction with doxycycline (*n* = 9 images with one image per technical replicate, three technical replicates per transduction, and three independent transductions per perturbation). Significance was determined using a two-sided Mann-Whitney *U*-test. Scale bar, 400 μm. **j,** Cell cycle phase of MDA-MB-231 cells after perturbation of endogenous *CCND1* and/or rescue with FLAG, FLAG-*CCND1*, or FLAG-*CCND1*(ΔExon 2) expression at 4 days after doxycycline-induction. The cell cycle phase was determined using flow cytometry. The bars denote the mean of three independent transductions and significance was determined using a two-sided Mann-Whitney *U*-test.

*CCND1* encodes cyclin D1, which promotes the G1/S phase transition during cell cycle progression by binding to cyclin-dependent kinases like *CDK6*. To confirm the importance of the *CCND1* isoform including exon 2, we assessed proliferation in human MDA-MB-231 cells after exon-specific Cas13 perturbation. Targeting either *CCND1* exon 2 or 3 with Cas13 inhibited proliferation approximately 2-fold in both 2D and 3D cultures (**Fig. 4d and Supplementary Fig. 4c-e**).

To understand how exon skipping impacts *CCND1* function, we used AlphaFold3^31^ to model the CCND1/CDK6 complex (**Fig. 4e and Supplementary Fig. 4f**). Residues encoded by exon 2 of *CCND1* are located at the binding interface, suggesting that the exon is important for complex formation and stabilization (**Fig. 4e**). As expected, the confidence of AlphaFold3-predicted CCND1(ΔExon 2)-CDK6 interactions was reduced, with a 2.4-fold decrease in the interface predicted template modeling (iPTM) score at the protein interface (**Supplementary Fig. 4f**). We hypothesized that skipping exon 2 would impair cell cycle progression by disrupting the interaction between CCND1 and CDK6. To test this experimentally, we overexpressed both *CCND1* isoforms, with and without exon 2. We then analyzed CDK6 co-purification after CCND1 immunoprecipitation (**Fig. 4f**). Overexpressing the full-length isoform resulted in *CDK6* co-purification, as expected. However, CCND1(ΔExon 2) completely abrogated CCND1-CDK6 complex formation (**Fig. 4f**).

Finally, we confirmed the functional importance of exon 2 inclusion for cell cycle progression. We expressed both *CCND1* isoforms while targeting the endogenous transcript with Cas13 (**Fig. 4g**). Importantly, we engineered multiple silent mutations in the full-length *CCND1* isoform to escape Cas13 knockdown (**Supplementary Fig. 4g**). Only the wild-type *CCND1* isoform restored cell proliferation and cell cycle progression in 2D and 3D cultures (**Fig. 4h, i**). Similarly, only wild-type *CCND1* restored normal cell cycling in MDA-MB-231 cells, showing a greater portion of cells in G1. In contrast, cells expressing *CCND1*(ΔExon 2) remained in G1 phase (**Fig. 4j and Supplementary Fig. 4h, i**). Altogether, *SF3B4* knockdown leads to impaired cell fitness and an accumulation of cells in G1 phase (**Fig. 2a and Supplementary Fig. 2d**). With CRISPore-seq, this phenotype is linked to a loss-of-function *CCND1* isoform, highlighting the importance of exon inclusion and isoform expression for cell fitness.

## Discussion

In this study, we introduced CRISPore-seq, a scalable CRISPR screening platform for isoform characterization via long-read sequencing, and, using it, examined how RBP perturbations affect full-length transcript structure. We identified changes in gene and isoform expression upon RBP perturbation. We linked transcriptional phenotypes to cell fitness phenotypes, uncovered distinct pathway signatures and evaluated the mechanisms of alternative splicing changes by integrating RBP binding data. We also related changes in isoform expression to functional consequences on cell proliferation using exon-specific RNA-targeting CRISPR perturbations. Specifically, we identified a *CCND1* isoform with a skipped exon as a potential driver of cell cycle arrest following *SF3B4* perturbation and validated its effects on cell proliferation and cell cycle progression. We also identified a potential mechanism of action by disrupting complex formation with CDK6 and demonstrated that exon skipping leads to an accumulation of cells that remain at the G1 checkpoint.

CRISPore-seq requires minimal specialized equipment and is easily scalable. It combines common droplet-based single-cell workflows with both short-read and long-read transcriptomics. This makes it readily available for widespread use. A key advantage of CRISPore-seq is its adaptability. It can be used effectively in many biological contexts, such as various disease models and diverse cell types. Additionally, its scalability can accommodate genome-scale perturbation studies and can target a range of genomic elements, from genes to noncoding variants.

For example, CRISPore-seq could dissect how noncoding genetic variants affect splicing. This includes single-nucleotide polymorphisms (SNPs) from genome-wide association studies (GWAS), which are often found in intronic regions^32,33^. By using CRISPR perturbations to target these SNPs, CRISPore-seq could monitor changes in aberrant splicing, providing direct mechanistic links between genotype and cellular phenotype (e.g. in T cells with autoimmune disease contexts)^34,35^. Establishing such links at scale has previously been challenging. Beyond its current capabilities, we also envision that further CRISPore-seq iterations could include the use of various CRISPR systems (e.g. CRISPR activation, base and prime editing, or RNA-targeting CRISPRs), the implementation of combinatorial or dose-dependent perturbations, and the incorporation of additional multi-omic readouts^36–42^. Given the tremendous benefit of long-read sequencing in distinguishing transcript isoforms in scRNA-seq, CRISPore-seq offers a scalable platform for understanding the transcript diversity resulting from genetic perturbations.

## Acknowledgments

We thank the entire Sanjana laboratory for their support and advice. We also thank Oxford Nanopore, the NYU Biology Genomics Core and the Zegar Family Foundation for sequencing resources. N.E.S. is supported by NIH/National Human Genome Research Institute (DP2HG010099, R01HG012790), NIH/National Cancer Institute (R01CA218668, R01CA279135), the NIH/National Institute of Allergy and Infectious Diseases (R01AI176601), the NIH/National Heart, Lung, and Blood Institute (NHLBI) (R01HL168247), the Simons Foundation for Autism Research (Genomics of ASD 896724), the MacMillan Center for the Study of the Noncoding Cancer Genome, and New York University and New York Genome Center funds. N.A. is supported by the Swedish Research Council (VR) international postdoc grant (2023-00494).

## Author contributions

S.M., A.S., and N.E.S. conceived the project. A.S., W.H.B., X.D., P.R., and Z.D. performed CRISPore-seq. S.M., N.A., R.E.Y., and A.S. analyzed the data. S.M. and R.E.Y. performed immunoprecipitation and phenotypic rescue experiments. S.M., R.E.Y, N.A., and N.E.S wrote the manuscript with input from all authors. N.E.S. and S.J. supervised the work.

## Declaration of interests

A.S. is a co-founder of TruEdit Bio and Osteologic Therapeutics. X.D., P.R., and S.J. are employees of and hold equity in Oxford Nanopore Technologies. N.E.S. is an adviser to Qiagen and a co-founder and adviser of TruEdit Bio and OverT Bio. The remaining authors declare no competing interests.

## Methods

### Cell culture

Wild-type MDA-MB-231 cell lines were acquired from American Type Culture Collection (HTB-26). *Rfx*Cas13d-NLS MDA-MB-231 cells were obtained from Liang et al.^44^. HEK293FT cells were obtained from Thermo Fisher Scientific (Thermo R70007). All cells were cultured in D10 medium: Dulbecco’s Modified Eagle Medium with high glucose and stabilized L-glutamine (Cytiva SH30022.01) supplemented with 10% Serum Plus II (Sigma-Aldrich 14009C) or HyClone FetalClone III serum (SH30109.03). Cells were maintained at 37°C with 5% CO2.

### CRISPore-seq: Library design

We constructed a library of 45 CRISPR inhibition (CRISPRi) guide RNAs (gRNAs) targeting 15 known splicing regulators and 6 non-targeting controls that do not target anywhere in the human genome (**Supplementary Table 1**). Guides that target the transcription start sites of these genes were selected from the Dolcetto-A CRISPR guide library^45^. Oligonucleotides were ordered and synthesized in arrayed format and cloned into lentiGuideFE-Puro (Addgene 52963)^46^.

A total of 5 µg of lentiGuideFE-Puro plasmid was digested with *Esp3*I (Thermo FD0454) for 30 min at 37°C. Digested plasmid was dephosphorylated using FastAP (Thermo EF0651) for 30 min at 37°C. The digest was then column-purified using the Zymo DNA Clean & Concentrator kit (D4014). Oligos for CRISPRi gRNAs were annealed and phosphorylated using T4 polynucleotide kinase (Thermo M0201L) using the following incubation protocol: 30 min at 37°C, 5 min at 95°C, drop down to 4°C at a ramp rate of 1°C/5 seconds. Annealed oligos were diluted with H_2_O 1:100 before ligated into the cut lentiGuideFE-Puro plasmid using T7 ligase (NEB M0318) for 10 min at room temperature. We used 2 µl of each ligation protocol to transform Stbl3 cells (NEB C3040I), incubated at room temperature for 2 min. Transformed cells were plated on LB agar plates containing 100 µg/ml ampicillin and incubated at 37°C for 16 hours. Individual colonies were picked and grown at 37°C with 220 rpm shaking overnight. Plasmids were purified using the QIAprep Spin Miniprep Kit (Qiagen 12123) and verified by Sanger sequencing (Genewiz). We then combined equimolar amounts of the gRNA plasmids to generate the pooled library. We performed short-read sequencing (Illumina MiSeq) to verify that all plasmids were present and measure the uniformity of their representation via the ratio of the 90^th^:10^th^ percentile plasmids. This uniformity ratio was 1.4.

### Lentivirus production

Lentiviral libraries were prepared in T225 flasks. Each flask was seeded with 27 × 10^6^ HEK293FT cells the day before in 30 ml of antibiotic-free D10 media to achieve 80-90% confluence before transfection. The transfection mix was 24.9μg of the transfer plasmid, 13.7 μg pMD2.G (Addgene 12260), 19.9 μg psPAX2 (Addgene 12259), 2490 μl OptiMEM (Invitrogen 51985-091) and 138 μl 1 mg/ml polyethylenimine linear MW 25000 (Polysciences 23966). The solution was mixed and incubated for 10 min at room temperature then added dropwise. After 5 hours, the media was changed to D10 with 1% bovine serum albumin (VWR AAJ65097). Viral supernatants were collected 72 hours following transfection, spun down, filtered with a 0.45mm filter (Millipore SE1M003M00) and stored at -80°C.

### Monoclonal CRISPRi cell line generation

Monoclonal CRISPRi-expressing cell lines were generated by transducing wild-type MDA-MB-231 cells with lentiCRISPRi(v2)-Blast (Addgene 170068)^46^ and selected with 5 μg/ml of blasticidin S (A.G. Scientific B-1247). Single-cell colonies were isolated by low-density plating and CRISPRi(v2) expression was confirmed via western blot.

### CRISPore-seq: Library preparation and sequencing

Pooled libraries were transduced into MDA-MB-231-CRISPRi cells, and, at 24 hours after transduction, cells were lifted and washed in PBS before being re-plated in puromycin-containing selection media. The survival was below 4% after 2 days of selection in 2 µg/mL puromycin (Invivogen ant-pr-1). On day 7 post-transduction, the cells were collected and processed for cell barcoding and transcriptome capture.

We performed cell-hashing and barcoding for ECCITE-seq^47^ and separated cells into 8 sub-pools. For each pool, we resuspended 1 million cells in 100 µl staining buffer (2% BSA, 0.01% Tween in PBS). We then added 10 µl Human TruStain FcX Receptor Blocking Solution (BioLegend) and incubated it on ice for 10 minutes. We then added 0.5 µg hashing antibodies to each of the 8 sub-pools (BioLegend), incubated on ice for 30 minutes and washed cells 3 times with 1 ml staining buffer. Cells were spun down at 4°C for 5 minutes at 450 ×g. Cells were resuspended in 50 µl ice cold PBS each and 1 µl was used to determine cell count. Cells from all 8 sub-pools were merged at equal proportions in 1 ml staining buffer, spun down at 350 ×g for 5 min at 4°C. Cells were resuspended in PBS at 1500 cell/µl and filtered through a 40 µm strainer (Flowmi).

Using the Chromium Single Cell Immune Profiling Solution v1.0 kit (10x Genomics), we targeted recovery of 12,000 cells per lane (superloading). We followed the 10x Genomics single cell assay with the following modifications: 5.9 µl of 2 µM gd_RT_V4 primer was added to the RT reaction and 1 µl of 1 µM of hashtag & guide-tag additive were added to the cDNA PCR as additives.

Post GEM-RT cleanup and cDNA amplification was carried out as described in the 10x Genomics single cell kit. Following cDNA library amplification, we set aside 300 ng of full-length cDNA library at 4° C for long-read sequencing. A total of 100 ng of cDNA was used to process the 5’ Gene Expression (GEX) library using fragmentation, end repair and A-tailing. Hashtag and guide-tag library preparations were performed as in CITE-seq^48^. Hashtag and guide RNA libraries were separated from cDNA libraries using 0.6× SPRI as desribed in Stoeckius *et al*. 2017 prior to fragmentation. Gene expression (GEX), hashtag and guide RNA libraries were quantified and diluted to 4 nmol and pooled in the following ratio: 5% Hashtag library, 10% Guide RNA library, 83% Gene expression library and 2% PhiX spike in. Each library was sequenced on a NextSeq550 75-cycle high-output run (Illumina).

A total of 300 ng of cDNA gene expression amplicon library was prepared for nanopore cDNA sequencing. No fragmentation was performed and full-length cDNA was quantified and sized using Qubit fluorometer and Agilent Bioanalyzer, respectively. The sequencing library was prepared using Oxford Nanopore Technologies’ SQK-LSK110 library preparation protocol and was sequenced on a PromethION FLO-PRO002 flowcell. Basecalling was performed with MinKNOW and Guppy v5.1.13 using the dna_r9.4.1_450bps_HAC model, yielding a total of 88 million cDNA reads.

### Cell proliferation and live imaging

To target specific exons for isoform perturbations, we used RNA-targeting *Rfx*Cas13d knockdown. First, we constructed a guide RNA (gRNA) expression plasmid (pLentiRNAGuide_005) by cloning a cassette encoding a human-codon optimized mStayGold-P2A-Puro from lentiGuideFE-mSG-Puro (Addgene 226522) into pLentiRNAGuide_001 (Addgene 138150). For this, we used *AgeI* and *ApaI* restriction sites and T7 ligase (New England Biolabs M0318L). Next, we cloned Cas13 gRNAs into pLentiRNAGuide_005 using *BsmBI* restriction sites and T7 ligation. These gRNAs targeted *CCND1* exon 2 (spacer: 5′-AGAUGCACAGCUUCUCGGCCGUC-3′), exon 3 (spacer: 5′-AACUUCACAUCUGUGGCACAGAG-3′), or were non-targeting controls (spacer: 5′-GAUAAGAGACUCUCGAUGUUGCG-3′). Then, we produced lentiviruses from these constructs.

For competitive growth assays, *Rfx*Cas13d MDA-MB-231 cells were transduced in triplicate. Cells were selected with 1 µg/ml puromycin for two days. Then, mStayGold-positive (perturbed) cells were co-cultured with parental MDA-MB-231 cells. They were mixed in equal ratios (1 × 10^5^ cells each per 24-well plate). After 24 hours, the initial proportion of mStayGold-positive cells was determined using an Incucyte S3 (Sartorius) at 20× magnification. Cas13 expression was then induced with 1 µg/ml doxycycline. The ratio of mStayGold-positive to mStayGold-negative cells was monitored for 6 days via live-cell imaging. Survival of perturbed cells was calculated by normalizing these ratios to the pre-induction baseline for each well. These values were then normalized to the median from co-cultures with non-targeting gRNA controls. Representative images show confluence masks (Incucyte Live Cell Analysis software).

For 2D and 3D proliferation assays, *Rfx*Cas13d MDA-MB-231 cells were first transduced in triplicate and selected with puromycin (1 µg/ml for 48 h). Then, 2 × 10^3^ cells were seeded into each well of 96-well plates and induced with doxycycline (1 µg/ml). Tissue culture-treated plates (Corning 3904) were used for 2D assays. For 3D assays, ultra-low attachment round-bottom plates (Corning 7007) were used and spheroid formation was induced by brief centrifugation (300 ×g for 5 min). Cell and spheroid proliferation were monitored using the Incucyte S3 system. 2D cultures were monitored for 4 days (10× magnification) and 3D spheroid cultures were monitored for 7 days (4× magnification). To calculate doubling times (*DT*), normalized cell confluence data (set to 1 at time = 0 h) was modeled using an exponential growth equation: confluence = exp(*k* × time), where *k* is the growth rate constant. This model was fitted to each perturbation-replicate group via non-linear least squares (`nls` in R v4.2), with an initial *k* = 0.05. The *DT* was computed as *DT* = ln(2)/*k*.

### Transgene rescue and cell cycle analysis

To rescue *Rfx*Cas13d-mediated *CCND1* perturbations, we first cloned overexpression cassettes into the pLentiRNAGuide_001 vector (Addgene 138150). We inserted PGK-driven coding sequences (expressing FLAG, FLAG-*CCND1*, and FLAG-*CCND1*(ΔExon 2)) using Gibson assembly with 25 ng of *NheI*-digested pLentiRNAGuide_001 and 100 ng of gBlocks (IDT) that included the respective expression cassette and overhangs for Gibson cloning. We inserted five silent mutations in the FLAG-*CCND1* sequence to ensure escape from *Rfx*Cas13d-mediated knockdown for rescue experiments when targeting the endogenous *CCND1* transcript (**Supplementary Fig. 4g**). Then, we cloned *Rfx*Cas13d gRNAs into these plasmids that target either the second exon of *CCND1* or do not target anywhere in the human transcriptome (non-targeting control) using *BsmBI* restriction sites and T7 ligation and produced lentiviruses.

For 2D and 3D proliferation assays, *Rfx*Cas13d MDA-MB-231 cells were first transduced with lentiviruses containing these gRNAs and overexpression cassettes, selected with puromycin (1 µg/ml for 48 h) and induced with doxycycline (1 µg/ml). We seeded 2 × 10^3^ cells into each well of 96-well plates (tissue culture-treated plates for 2D assays, ultra-low attachment round-bottom plates for 3D assays). Cell and spheroid proliferation were monitored using the Incucyte S3 system and analyzed using the Incucyte Live Cell Analysis software.

For cell cycle assays, we used propidium iodide (PI) staining and flow cytometry. 5 × 10^5^ cells per condition were pelleted in 15-ml tubes, washed once with 1 ml of ice-cold PBS and resuspended in 1 ml of ice-cold PBS. Cells were fixed with an additional 3 ml of ice-cold 100% ethanol to a final concentration of 75% overnight. Cells were then spun down at 500 ×g for 5 min, washed with 1 ml PBS and stained with 250 μl of FxCycle PI/RNAse solution (Thermo F10797) for 30 min at room temperature in the dark for flow cytometry (Sony SH800). Gating was performed with exclusion of debris on the basis of FSC-A/SSC-H, followed by gating on singlets with FSC-A/FSC-H. The cell-cycle profile was modeled on the PI signal intensity using FlowJo (version 10.10.0) with a Dean-Jett Fox model.

### Immunoprecipitation

We transfected 5 × 10^6^ HEK293FT cells with expression vectors for FLAG-tagged cyclin D1 variants under a constitutive PGK promoter using Lipofectamine 3000 (Thermo L3000008). Transfected cells were selected for 48 hours using 1 µg/ml puromycin. After that, 1 × 10^7^ cells were collected, washed with 1× DPBS (Sigma D8537) and lysed with 1 ml of RIP buffer (50 mM Tris-HCl buffer pH 7.5 (Thermo 15567027), 100 mM sodium chloride (Sigma, S5150), 1% NP-40 (Thermo 85124), 0.1% SDS (Sigma, L3771) and 0.5% sodium deoxycholate (Sigma, D6750)). Immediately before using, we added 5 µl of benzonase (Sigma, E8263) per 1 ml of RIP buffer and performed lysis for 15 min at 4°C. Samples were then spun down for 10 min at 4°C and 50 µl of the lysate was set aside as an input control.

Dynabeads Protein A Beads (Thermo 10003D) (50 µl per sample) were washed with RIP buffer 3 times and incubated with 5 µg per sample mouse anti-DYKDDDDK tag monoclonal antibody (Proteintech 66008-4-Ig) for 30 min at room temperature. Antibody-bead complexes were then incubated with the remaining cell lysate for 2 hours at 4°C. Protein enrichment compared to input samples was analyzed by western blotting, as described below.

### Western blot

Samples were denatured in NuPAGE LDS Sample Buffer (Thermo P0007) supplemented with 100 mM DTT (Cayman 700416) for 10 min at 65°C. Denatured samples and PageRuler prestained protein ladder (Thermo 26616) were separated in Novex 4-12% Tris-Glycine mini gels (Thermo XP04125) in 1× MOPS running buffer (Thermo NP0001) for 1 hour at 160 V, then transferred to nitrocellulose membranes (BioRad 1620112) in 1x Tris-Glycine transfer buffer (Thermo LC3675) supplemented with 10% methanol for 1 hour at 30 V.

Membranes were blocked in 5% milk (Research Products International M17200) dissolved in 1× TBS with 0.1% Tween-20 (TBS-T) for 60 min. The membranes were then incubated overnight at 4°C with the following primary antibodies: Anti-DYKDDDDK (FLAG) tag mouse monoclonal antibody (Proteintech 66008-4-Ig), CDK6 monoclonal antibody (Proteintech 66278-1-Ig), or Beta Actin Recombinant antibody (Proteintech 81115-1-RR). All primary antibodies were used at a 1:2,000 dilution in 5% BSA (VWR, AAJ65097) dissolved in 1× TBS-T. The blots were washed and then incubated with IRDye 800CW donkey anti-rabbit or anti-mouse secondary antibodies (LI-COR 925-32213 or LI-COR 925-32212, 1:10,000 dilution in 5% BSA/TBS-T) for 60 min at room temperature, washed three times with TBS-T and visualized using an Odyssey CLx imaging system (LI-COR).

### CRISPore-seq: Alignment and splicing quantification

Short-read data was aligned using CellRanger v7.1.0 using GRCh38 (http://ftp.ensembl.org/pub/release-109/fasta/homo_sapiens/dna/Homo_sapiens.GRCh38.dna.primary_assembly.fa.gz) with Gencode v47 annotations (http://ftp.ebi.ac.uk/pub/databases/gencode/Gencode_human/release_47/gencode.v47.primary_as sembly.annotation.gtf.gz). Long-read data was processed into gene and isoform-specific count-matrices using Oxford Nanopore Technologies’ single-cell workflow (wf-single-cell, https://github.com/epi2me-labs/wf-single-cell) using the Gencode v47 genome annotation, with the following flags: `--expected_cells 12000` and `--kit ’5prime:v2’`.

To compare the coverage between short-read and long-read sequencing, the per-nucleotide coverage was calculated using BEDTools (v2.31.0) to compute read depth at each base position across all exons for ten highly-expressed transcripts: *GAPDH (ENST00000229239*), *ACTB (ENST00000646664)*, *RPL12 (ENST00000361436)*, *FTL (ENST00000331825)*, *RPLP0 (ENST00000392514)*, *RPL7A (ENST00000323345)*, *RPL18 (ENST00000549920*), *MT2A (ENST00000245185*), *RPL30 (ENST00000287038)*, *PFN1 (ENST00000225655)*. For each transcript, genomic positions were normalized to percent distance along the transcript body (0-100%). Coverage values were normalized per transcript.

Aligned data were analyzed using Seurat v5.0^49^, SingleCellExperiment v1.20.1^50^, and SCEPTRE v0.3.0^51^. Cells with a low number of features (<400 for long-read data and <3000 for short-read data) were filtered out as low-quality cells. Guide RNA expression was determined from short-read data, assigning to each cell the guide RNA with the maximum count in that cell. For the single-cell expression heatmap of downregulated protein-coding genes, we used the `FindMarkers` function, comparing each perturbation to the non-targeting control. We then randomly sampled 50 cells per group and used the `NormalizeData` and `ScaleData` functions before heatmap visualization. To assign the cell cycle phase of individual cells, we used the `CellCycleScoring` function, which calculates scores for G1, S and G2/M phases based on the expression of predefined marker genes. For low-dimensional embedding of perturbation effects, the top 2000 variable features with the highest variance relative to their mean expression were used for multidimensional scaling analysis. Pseudocells were generated by averaging expression profiles of 50 randomly sampled cells per condition (1,000 pseudocells per perturbation). Euclidean distances were calculated between all pseudocells, and classical MDS embedding was performed using the `cmdscale` function in R. Differentially-expressed genes were identified using SCEPTRE (*P*_Adj._ < 0.05, Benjamini-Hochberg adjusted, **Supplementary Table 2**)^51^.

For alternative splicing analysis, scNanoGPS v1.1^52^ was used to pseudo-bulk transcripts by perturbation. The `scanner` module of scNanoGPS was used to process raw sequencing reads and detect barcodes. As scNanoGPS is designed to use 3′ Chromium kits certain input parameters were adjusted to process 5′ data. The input 5′ and 3′ sequences were swapped, and the poly(A) sequence was replaced with the template switch oligo (TSO) sequence using the following flags: `--a3=CTACACGACGCTCTTCCGATCT --a5=AAGCAGTGGTATCAACGCAGAGTAC --pT=TTTCTTATATGGG --lUMI=10`

We then removed the TSO sequence from FASTQ files generated by the scanner module using Biopython’s `pairwise2` module to perform local sequence alignment, searching for the TSO sequence within the first 40 bases of each read. Reads containing a match above a defined threshold were trimmed to remove the TSO sequence. After barcode detection and TSO removal, the `assigner` module of scNanoGPS was used to assign cell barcodes to sequencing reads. Finally, the `curator` module of scNanoGPS was used to process and refine cell barcode assignments, using GRCh38 as the reference genome. The curated reads were merged by perturbation target and used for alternative splicing analysis.

FLAIR v2.0^53^ was used to identify alternative splicing events associated with specific perturbations. The FLAIR `align` module mapped merged FASTQ files to the GRCh38 reference genome. The FLAIR `correct` module was used to refine misaligned splice sites using the Gencode v47 annotation, adjusting junctions to known transcript structures for improved accuracy. FLAIR `collapse` was then used to generate a high-confidence isoform set by filtering redundant transcripts, enforcing criteria (`--stringent`), verifying splice junctions (`--check_splice`), and prioritizing annotated isoforms (`--annotation_reliant`), as recommended. Finally, FLAIR `quantify` estimated transcript abundance by mapping reads to the collapsed isoforms, and FLAIR `diffSplice` and `diffsplice_fishers_exact` were used to identify alternative splicing events associated with each perturbation. Events with a change in percent spliced-in |ΔΨ| > 0.05 and *P* < 0.05 were identified as significant AS events.

### Gene set enrichment analyses

Gene set enrichment analyses (GSEA) were performed on pre-ranked lists using the R package clusterProfiler (v4.10.0) and MSigDB pathways (v2024.1)^54,55^. We used gene set files from Gene Ontology (Biological processes, c5.go.bp.v2024.1.Hs.symbols.gmt)^56^ and Reactome (c2.cp.reactome.v2024.1.Hs.symbols.gmt)^57^ databases.

The `GSEA` function was used with the following key parameters: `TERM2GENE = geneSetFile`, `exponent = 0`, `pAdjustMethod = “FDR”`, `pvalueCutoff = 1`, `by = “fgsea”`. As input, protein-coding genes were ranked by their log_2_ fold change after RBP perturbations, as determined from both short- and long-read CRISPore-seq data. *P* values from Kolmogorov-Smirnov tests were adjusted for multiple comparisons using the Benjamini-Hochberg method.

To find the overlap of significantly regulated Gene Ontology (GO) and Reactome pathways (*P*_Adj._ < 0.05) among different RBP perturbations, GSEA results were classified in significantly up- (normalized enrichment score [NES] > 0) or downregulated (NES < 0) pathways for each RBP. The number of RBPs commonly regulating each pathway was then quantified and categorized (e.g. pathway regulated by 1 RBP, 2 RBPs, 3-5 RBPs, 6-10 RBPs and 11-15 RBPs) (**Supplementary Table 3**). These pathways overlap statistics were visualized using the circlize R package (v0.4.16).

### eCLIP enrichment analysis

Enhanced Crosslinking and Immunoprecipitation (eCLIP) peaks from ENCODE (https://www.encodeproject.org) were used to define RBP binding sites^58,59^. Peak files containing enrichment values over size-matched input were obtained from ENCODE and peaks were considered significant if they showed at least a log_2_ fold enrichment of 2 over size-matched input and a *P* value < 0.001. Peaks from both K562 and HepG2 cell lines were included in the analysis. For RBPs with multiple replicates, peaks meeting the significance criteria in any replicate were considered valid binding sites. Each peak was assigned to its overlapping gene using Gencode v47 gene annotations.

eCLIP peak data were downloaded as bigBed files (**Supplementary Table 5**). The rtracklayer package (version 1.64.0) was used to import bigBed files into GRanges objects. For each alternative splicing event, genomic coordinates were parsed, and strand-specific upstream and downstream windows were defined using functions from the GenomicRanges (version 1.56.0) and IRanges (version 2.38.0) packages. Upstream windows were defined as -300 bp to +50 bp relative to the 3′ splice site, and downstream windows as -50 bp to +300 bp relative to the 5′ splice site of the alternative exon. For intron retention events, windows were defined as 100 bp upstream of the intron start, the intron itself, and 100 bp downstream of the intron end. The `findOverlaps` function from GenomicRanges was used to determine if any RBP-specific eCLIP peaks overlapped with the defined genomic windows for each splicing event. To visualize the positional enrichment of RBP binding sites relative to regulated splice sites, we generated binding heatmaps. For exon skipping events, regions flanking the 3′ splice site (−300 bp to +50 bp) and 5′ splice site (−50 bp to +300 bp) of the alternative exon were divided into consecutive 10 bp bins. For each RBP, we calculated the fraction of alternative splicing events that showed eCLIP peak overlaps within each 10 bp bin.

For the visualization of eCLIP tracks in specific transcripts, the unique and strand-specific signals were downloaded as bigwig files (**Supplementary Table 5**). The read coverage from two independent eCLIP replicates per RBP and the corresponding size-matched input was visualized using the Integrative Genomics Viewer (IGV, https://igv.org).

### Visualization of alternative splicing events

For visualization and analysis of exon-specific splicing events, we developed a custom analysis pipeline that processes BAM files to examine read coverage and exon inclusion patterns across perturbations. The pipeline was implemented using Python 3.8 and R (tidyverse v2.0.0), generating normalized coverage plots and exon inclusion rates for comparative analysis between perturbation and control conditions. Gene-specific regions were extracted from aligned BAM files using samtools (v1.15) with Gencode v47 gene annotations. Reads were then categorized based on inclusion or exclusion of the region of interest. Secondary alignments and reads with mapping quality below 20 were excluded from the analysis. Coverage was calculated at single-nucleotide resolution using samtools depth and normalized to maximum coverage within each condition. For visualizing genes with large introns, intronic regions with coverage below 1 read were truncated, while maintaining a 200-base-pair window at gap boundaries to preserve context at exon-intron junctions.

### Prediction of protein structures

Wild-type amino-acid sequences were obtained from UniProt. Exon-skipped isoforms were generated using IGV to identify amino acid sequences of isoforms with alternative splicing events. Structural predictions and interaction models were generated using the AlphaFold Server (AlphaFold 3)^60^. For each query, only the top-ranked prediction (based on AlphaFold’s internal ranking score) was used. Model confidence was assessed using the predicted template modeling score (pTM) for global structure and the interface pTM (ipTM) for inter-chain interactions. 3D structural renderings were visualized using Mol* Viewer^61^.

## Materials availability

Further information and requests for resources and reagents should be directed to and will be fulfilled by the Lead Contact, Neville Sanjana.

## Data and code availability

Genomic datasets are available via <u>BioProject</u> (PRJNA1369506). Gene annotations were downloaded from <u>GENCODE</u> (v47) and the reference genome from <u>ENSEMBL</u> (GRCh38). RNA-binding protein eCLIP data were downloaded from <u>ENCODE</u>.

## Supplementary Tables

**Supplementary Table 1:** CRISPore-seq sgRNA library targeting RNA-binding proteins.

**Supplementary Table 2:** Differential gene expression upon RNA-binding protein perturbations using short-read scRNA-seq.

**Supplementary Table 3:** Gene set enrichment analyses upon RNA-binding protein perturbations.

**Supplementary Table 4:** Alternative splicing events identified in CRISPore-seq.

**Supplementary Table 5:** Enhanced Crosslinking and Immunoprecipitation (eCLIP) ENCODE data for RNA-binding proteins.

**Supplementary Table 6:** Binding of RNA-binding proteins in alternatively spliced exons after perturbation.

**Supplementary Table 7:** Binding of RNA-binding proteins in alternatively spliced introns after perturbation.

**Supplementary Table 8:** Exon-specific Cas13 guide RNA depletion in *Rfx*Cas13d MDA-MB-231 cells.

## Notes

### Competing Interest Statement

A.S. is a co-founder of TruEdit Bio and Osteologic Therapeutics. D.D., P.R., and S.J. are employees of and hold equity in Oxford Nanopore Technologies. N.E.S. is an adviser to Qiagen and a co-founder and adviser of TruEdit Bio and OverT Bio. The remaining authors declare no competing interests.

## References

1. Berk, A. J. & Sharp, P. A. Sizing and mapping of early adenovirus mRNAs by gel electrophoresis of S1 endonuclease-digested hybrids. Cell 12, 721–732 (1977).

2. Chow, L. T., Roberts, J. M., Lewis, J. B. & Broker, T. R. A map of cytoplasmic RNA transcripts from lytic adenovirus type 2, determined by electron microscopy of RNA:DNA hybrids. Cell 11, 819–836 (1977).

3. Rogalska, M. E., Vivori, C. & Valcárcel, J. Regulation of pre-mRNA splicing: roles in physiology and disease, and therapeutic prospects. Nat Rev Genet 24, 251–269 (2023).

4. Marasco, L. E. & Kornblihtt, A. R. The physiology of alternative splicing. Nat Rev Mol Cell Biol 24, 242–254 (2023).

5. Pan, Q., Shai, O., Lee, L. J., Frey, B. J. & Blencowe, B. J. Deep surveying of alternative splicing complexity in the human transcriptome by high-throughput sequencing. Nat Genet 40, 1413–1415 (2008).

6. Wang, E. T. et al. Alternative isoform regulation in human tissue transcriptomes. Nature 456, 470–476 (2008).

7. Nilsen, T. W. & Graveley, B. R. Expansion of the eukaryotic proteome by alternative splicing. Nature 463, 457–463 (2010).

8. Barbosa-Morais, N. L. et al. The Evolutionary Landscape of Alternative Splicing in Vertebrate Species. Science 338, 1587–1593 (2012).

9. Merkin, J., Russell, C., Chen, P. & Burge, C. B. Evolutionary Dynamics of Gene and Isoform Regulation in Mammalian Tissues. Science 338, 1593–1599 (2012).

10. Bradley, R. K. & Anczuków, O. RNA splicing dysregulation and the hallmarks of cancer. Nat Rev Cancer 23, 135–155 (2023).

11. Nikom, D. & Zheng, S. Alternative splicing in neurodegenerative disease and the promise of RNA therapies. Nat Rev Neurosci 24, 457–473 (2023).

12. Van Nostrand, E. L. et al. A large-scale binding and functional map of human RNA-binding proteins. Nature 583, 711–719 (2020).

13. Rogalska, M. E. et al. Transcriptome-wide splicing network reveals specialized regulatory functions of the core spliceosome. Science 386, 551–560 (2024).

14. Shalem, O. et al. Genome-scale CRISPR-Cas9 knockout screening in human cells. Science 343, 84–87 (2014).

15. Wang, T., Wei, J. J., Sabatini, D. M. & Lander, E. S. Genetic screens in human cells using the CRISPR-Cas9 system. Science 343, 80–84 (2014).

16. Hart, T. et al. High-Resolution CRISPR Screens Reveal Fitness Genes and Genotype-Specific Cancer Liabilities. Cell 163, 1515–1526 (2015).

17. Ghandi, M. et al. Next-generation characterization of the Cancer Cell Line Encyclopedia. Nature 569, 503–508 (2019).

18. Dixit, A. et al. Perturb-Seq: Dissecting Molecular Circuits with Scalable Single-Cell RNA Profiling of Pooled Genetic Screens. Cell 167, 1853–1866.e17 (2016).

19. Jin, X. et al. In vivo Perturb-Seq reveals neuronal and glial abnormalities associated with autism risk genes. Science 370, eaaz6063 (2020).

20. Replogle, J. M. et al. Mapping information-rich genotype-phenotype landscapes with genome-scale Perturb-seq. Cell 185, 2559–2575.e28 (2022).

21. Morris, J. A. et al. Discovery of target genes and pathways at GWAS loci by pooled single-cell CRISPR screens. Science 380, eadh7699 (2023).

22. Mimitou, E. P. et al. Multiplexed detection of proteins, transcriptomes, clonotypes and CRISPR perturbations in single cells. Nat Methods 16, 409–412 (2019).

23. Barry, T., Mason, K., Roeder, K. & Katsevich, E. Robust differential expression testing for single-cell CRISPR screens at low multiplicity of infection. Genome Biol 25, 124 (2024).

24. Gerstberger, S., Hafner, M. & Tuschl, T. A census of human RNA-binding proteins. Nat Rev Genet 15, 829–845 (2014).

25. Hentze, M. W., Castello, A., Schwarzl, T. & Preiss, T. A brave new world of RNA-binding proteins. Nat Rev Mol Cell Biol 19, 327–341 (2018).

26. Tsherniak, A. et al. Defining a Cancer Dependency Map. Cell 170, 564–576.e16 (2017).

27. Ashburner, M. et al. Gene Ontology: tool for the unification of biology. Nat Genet 25, 25–29 (2000).

28. Milacic, M. et al. The Reactome Pathway Knowledgebase 2024. Nucleic Acids Res 52, D672–D678 (2024).

29. Van Nostrand, E. L. et al. Robust transcriptome-wide discovery of RNA-binding protein binding sites with enhanced CLIP (eCLIP). Nat Methods 13, 508–514 (2016).

30. Liang, W.-W., et al. Transcriptome-scale RNA-targeting CRISPR screens reveal essential lncRNAs in human cells. *Figshare preprint* 10.6084/M9.FIGSHARE.30370171.V1 (2025) doi:10.6084/M9.FIGSHARE.30370171.V1.

31. Abramson, J. et al. Accurate structure prediction of biomolecular interactions with AlphaFold 3. Nature 630, 493–500 (2024).

32. Li, Y. I. et al. RNA splicing is a primary link between genetic variation and disease. Science 352, 600–604 (2016).

33. Qi, T. et al. Genetic control of RNA splicing and its distinct role in complex trait variation. Nat Genet 54, 1355–1363 (2022).

34. Farh, K. K.-H. et al. Genetic and epigenetic fine mapping of causal autoimmune disease variants. Nature 518, 337–343 (2015).

35. Harroud, A. & Hafler, D. A. Common genetic factors among autoimmune diseases. Science 380, 485–490 (2023).

36. Schmidt, R. et al. CRISPR activation and interference screens decode stimulation responses in primary human T cells. Science 375, eabj4008 (2022).

37. Komor, A. C., Kim, Y. B., Packer, M. S., Zuris, J. A. & Liu, D. R. Programmable editing of a target base in genomic DNA without double-stranded DNA cleavage. Nature 533, 420–424 (2016).

38. Anzalone, A. V. et al. Search-and-replace genome editing without double-strand breaks or donor DNA. Nature 576, 149–157 (2019).

39. Wessels, H.-H. et al. Efficient combinatorial targeting of RNA transcripts in single cells with Cas13 RNA Perturb-seq. Nat Methods 20, 86–94 (2023).

40. Hart, S. K. et al. Precise RNA targeting with CRISPR-Cas13d. Nat Biotechnol 10.1038/s41587-025-02558-3 (2025) doi:10.1038/s41587-025-02558-3.

41. Rubin, A. J. et al. Coupled Single-Cell CRISPR Screening and Epigenomic Profiling Reveals Causal Gene Regulatory Networks. Cell 176, 361–376.e17 (2019).

42. Yan, R. E. et al. Pooled CRISPR screens with joint single-nucleus chromatin accessibility and transcriptome profiling. Nat Biotechnol 10.1038/s41587-024-02475-x (2024) doi:10.1038/s41587-024-02475-x.

43. Liang, W.-W. et al. Transcriptome-scale RNA-targeting CRISPR screens reveal essential lncRNAs in human cells. Cell 187, 7637–7654.e29 (2024).

44. Liang, W.-W. et al. Transcriptome-scale RNA-targeting CRISPR screens reveal essential lncRNAs in human cells. Cell 187, 7637–7654.e29 (2024).

45. Sanson, K. R. et al. Optimized libraries for CRISPR-Cas9 genetic screens with multiple modalities. Nat Commun 9, 5416 (2018).

46. Morris, J. A. et al. Discovery of target genes and pathways at GWAS loci by pooled single-cell CRISPR screens. Science 380, eadh7699 (2023).

47. Mimitou, E. P. et al. Multiplexed detection of proteins, transcriptomes, clonotypes and CRISPR perturbations in single cells. Nat Methods 16, 409–412 (2019).

48. Stoeckius, M. et al. Simultaneous epitope and transcriptome measurement in single cells. Nat Methods 14, 865–868 (2017).

49. Hao, Y. et al. Dictionary learning for integrative, multimodal and scalable single-cell analysis. Nat Biotechnol 42, 293–304 (2024).

50. Amezquita, R. A. et al. Orchestrating single-cell analysis with Bioconductor. Nat Methods 17, 137–145 (2020).

51. Barry, T., Mason, K., Roeder, K. & Katsevich, E. Robust differential expression testing for single-cell CRISPR screens at low multiplicity of infection. Genome Biol 25, 124 (2024).

52. Shiau, C.-K. et al. High throughput single cell long-read sequencing analyses of same-cell genotypes and phenotypes in human tumors. Nat Commun 14, 4124 (2023).

53. Tang, A. D. et al. Full-length transcript characterization of SF3B1 mutation in chronic lymphocytic leukemia reveals downregulation of retained introns. Nat Commun 11, 1438 (2020).

54. Subramanian, A. et al. Gene set enrichment analysis: a knowledge-based approach for interpreting genome-wide expression profiles. Proc Natl Acad Sci U S A 102, 15545–15550 (2005).

55. Liberzon, A. et al. The Molecular Signatures Database (MSigDB) hallmark gene set collection. Cell Syst 1, 417–425 (2015).

56. Ashburner, M. et al. Gene Ontology: tool for the unification of biology. Nat Genet 25, 25–29 (2000).

57. Milacic, M. et al. The Reactome Pathway Knowledgebase 2024. Nucleic Acids Res 52, D672–D678 (2024).

58. Van Nostrand, E. L. et al. Robust transcriptome-wide discovery of RNA-binding protein binding sites with enhanced CLIP (eCLIP). Nat Methods 13, 508–514 (2016).

59. Van Nostrand, E. L. et al. A large-scale binding and functional map of human RNA-binding proteins. Nature 583, 711–719 (2020).

60. Abramson, J. et al. Accurate structure prediction of biomolecular interactions with AlphaFold 3. Nature 630, 493–500 (2024).

61. Sehnal, D. et al. Mol* Viewer: modern web app for 3D visualization and analysis of large biomolecular structures. Nucleic Acids Res 49, W431–W437 (2021).

