## Supplementary Figures for "Transcriptome-wide profiling of alternative splicing regulators with CRISPore-seq"

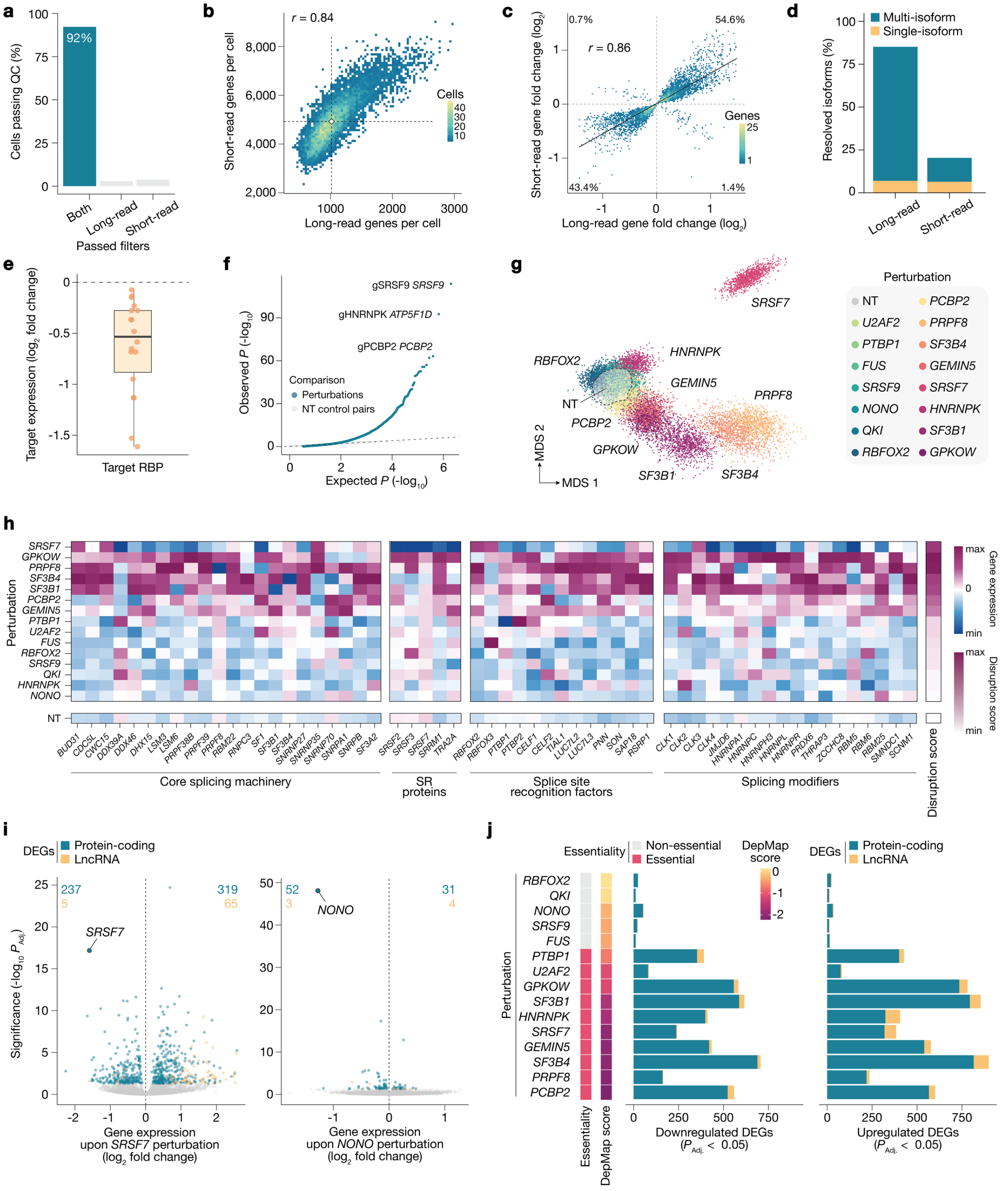

**Supplementary Fig. 1: Pooled CRISPR screens with joint short- and long-read transcriptomics.**

**a,** Percentage of cells passing filters for long-read and short-read sequencing (see *Methods*). Cells passing both filters were considered in downstream analysis. **b,** Pearson correlation between genes detected per cell as measured by short- and long-read sequencing. **c,** Gene expression changes as measured by short- and long-read sequencing with fold-change over control (non-targeting) cells. **d,** Resolved isoforms using short- and long-read sequencing. **e,** Pseudobulk expression of RNA-binding proteins (RBPs) targeted by CRISPR inhibition relative to non-targeting controls by perturbation. Boxplot shows median, interquartile range, and whiskers indicating 1.5× interquartile range. **f,** Quantile-quantile plot of gene expression changes induced by perturbations (*blue*) calculated with SCEPTRE[1], as well as non-targeting (NT) control pairs (*gray*). Dotted line indicates expected distribution. **g,** Multidimensional scaling (MDS) of cell transcriptomes from short-read data, colored by CRISPR perturbations. Each MDS point represents average expression of 50 cells per perturbation ($n$ = 1000 samples with replacement). **h,** Heatmap of expression changes (short-read) with affected genes grouped by role in spliceosome machinery. **i,** Differential expression of protein-coding genes and lncRNAs after knockdown of *SRSF7* and *NONO.* **j,** Number and type of differentially expressed genes (DEGs) after RBP perturbation from CRISPore-seq short-read sequencing and RBP essentiality in MDA-MB-231 cells from the Cancer Dependency Map (DepMap)[2].

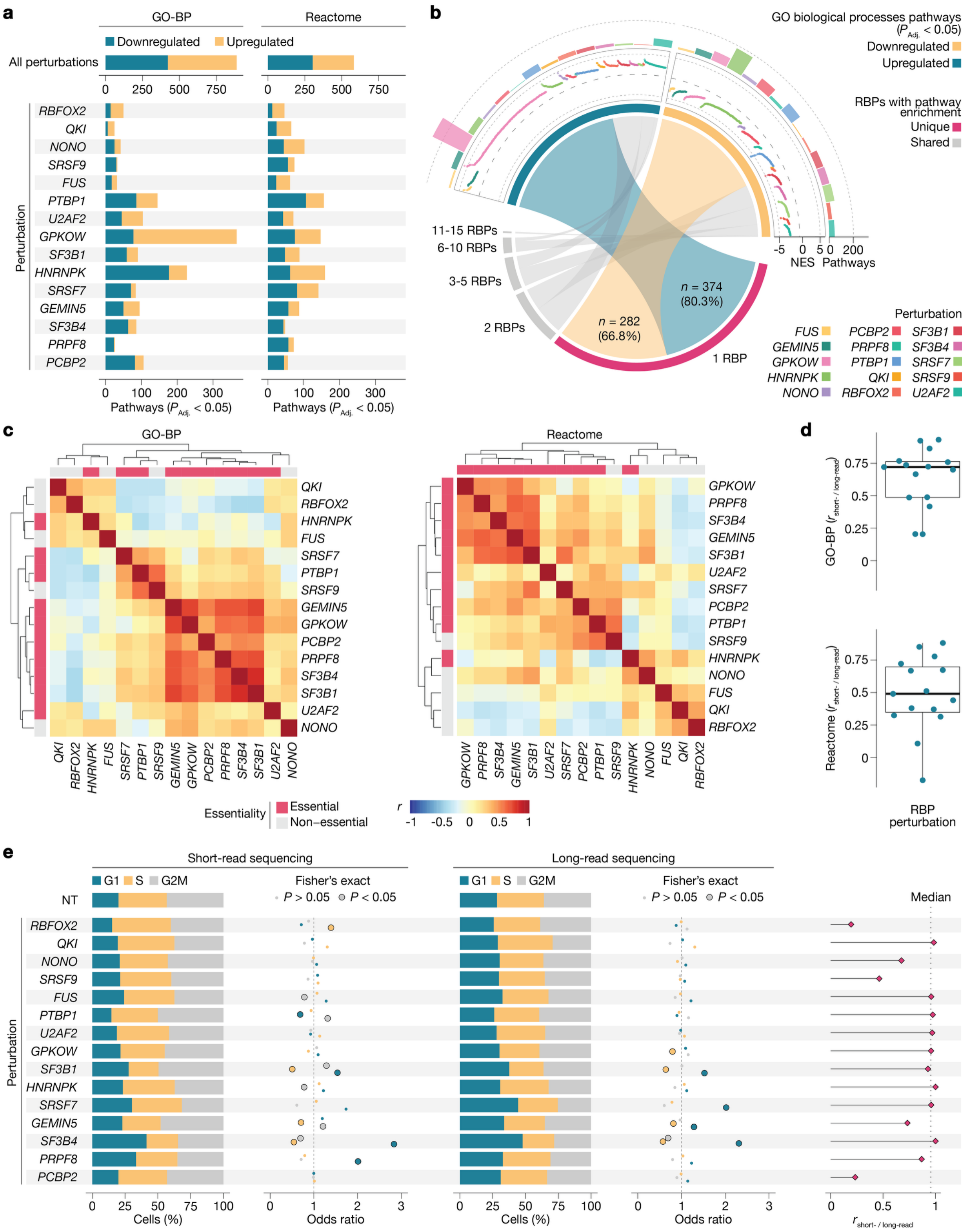


**Supplementary Fig. 2: Transcriptional phenotypes after RNA-binding protein perturbations.**

**a,** Number of significantly affected pathways ($P_{adj.} < 0.05$) from gene set enrichment analyses (GSEA) upon RNA-binding protein (RBP) perturbations using two curated pathway annotations from the Gene Ontology (GO) and Reactome databases[3,4]. **b,** Altered Gene Ontology (GO) pathways in cells with RBP perturbations. Outer tracks show the number of significant pathways per RBP ($P_{adj.} < 0.05$) and normalized enrichment scores (NES). The central chord diagram shows pathway overlaps among RBP perturbations. Pathway significance was assessed by a Kolmogorov-Smirnov test (Benjamini-Hochberg corrected). **c,** Pearson correlation heatmap for significant pathways ($P_{Adj.} < 0.05$) from GSEA analyses among RBP perturbations and RBP essentiality in MDA-MB-231 cells from the Cancer Dependency Map (DepMap)[2]. **d,** Pearson correlation for significant pathways ($P_{Adj.} < 0.05$) comparing short-read versus long-read sequencing data. **e,** Cell cycle phase distributions of individual perturbed cells assigned using CRISPore-seq short- (*left*) and long-read (*middle*) sequencing. The odds ratio and significance were computed by comparing the respective cell cycle phase distribution in perturbed cells to the fraction of cells that received non-targeting (NT) controls (Fisher's exact test). Pearson correlation of odds ratios for cell cycle phase enrichment between short- and long-read data (*right*).

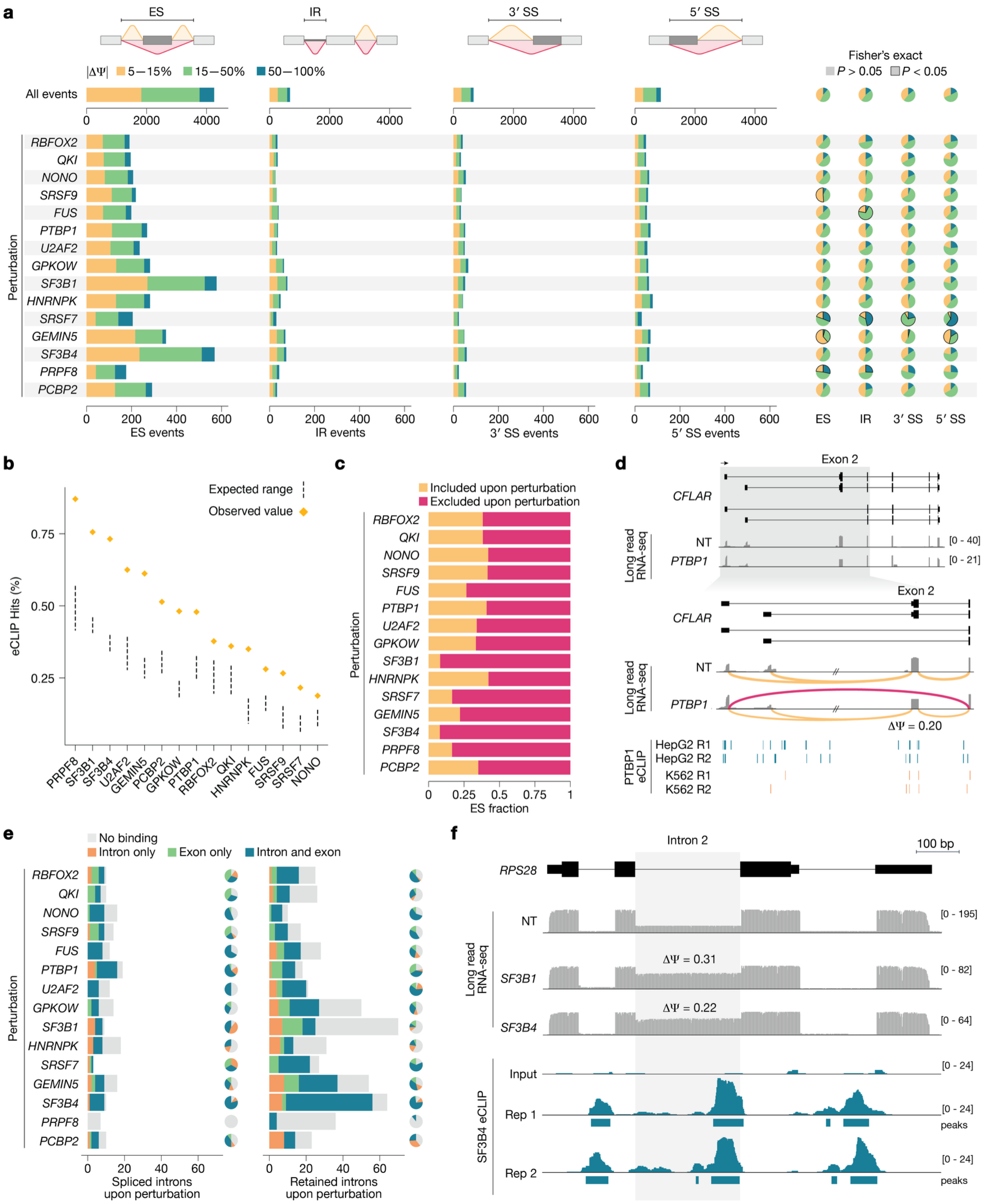


**Supplementary Fig. 3: Alternative splicing across RNA-binding protein perturbations.**
**a,** Number of total alternative splicing (AS) events ($P < 0.05$) of each class, including exon skipping (ES), intron retention (IR) and alternative 3′/5′ splice sites (SS). Splicing events are grouped by their magnitude in percent spliced-in $\Delta\Psi$ per AS type: strong if $|\Delta\Psi| > 50\%$, moderate if $15\% < |\Delta\Psi| < 50\%$, weak if $5\% < |\Delta\Psi| < 15\%$. Significant enrichment of splicing magnitude

groups was computed by comparing the respective fraction per perturbation to the fraction across all AS events using Fisher's exact test. **b,** Enrichment of enhanced Crosslinking and Immunoprecipitation (eCLIP) binding targets among genes with alternative splicing events. Points show the observed proportion of AS-affected genes that are direct RBP targets based on eCLIP data. Dashed lines indicate the expected proportion ± standard deviation from random sampling of protein-coding genes ($n$ = 100 iterations). **c,** Fraction of significant exon skipping events for exons that are included or excluded upon RBP perturbation. **d,** *CFLAR* splicing changes after *PTBP1* knockdown from long-read sequencing and eCLIP data showing *PTBP1* binding sites. Loops represent the relative frequency of exon inclusion (*yellow*) or exclusion (*pink*). **e,** Number of intron retention events, including retained introns and spliced introns after RNA-binding protein (RBP) perturbation, from CRISPore-seq long-read transcriptomics. Direct RBP binding was determined within the alternatively spliced intron or flanking exons (upstream: -100 nt, downstream: +100 nt) using RBP-specific eCLIP data. **f,** *RPS28* intron retention from long-read scRNA-seq after knockdown of *SF3B1* or *SF3B4*. Tracks below show the eCLIP read coverage and significant peaks for *SF3B4*-binding within the retained intron.

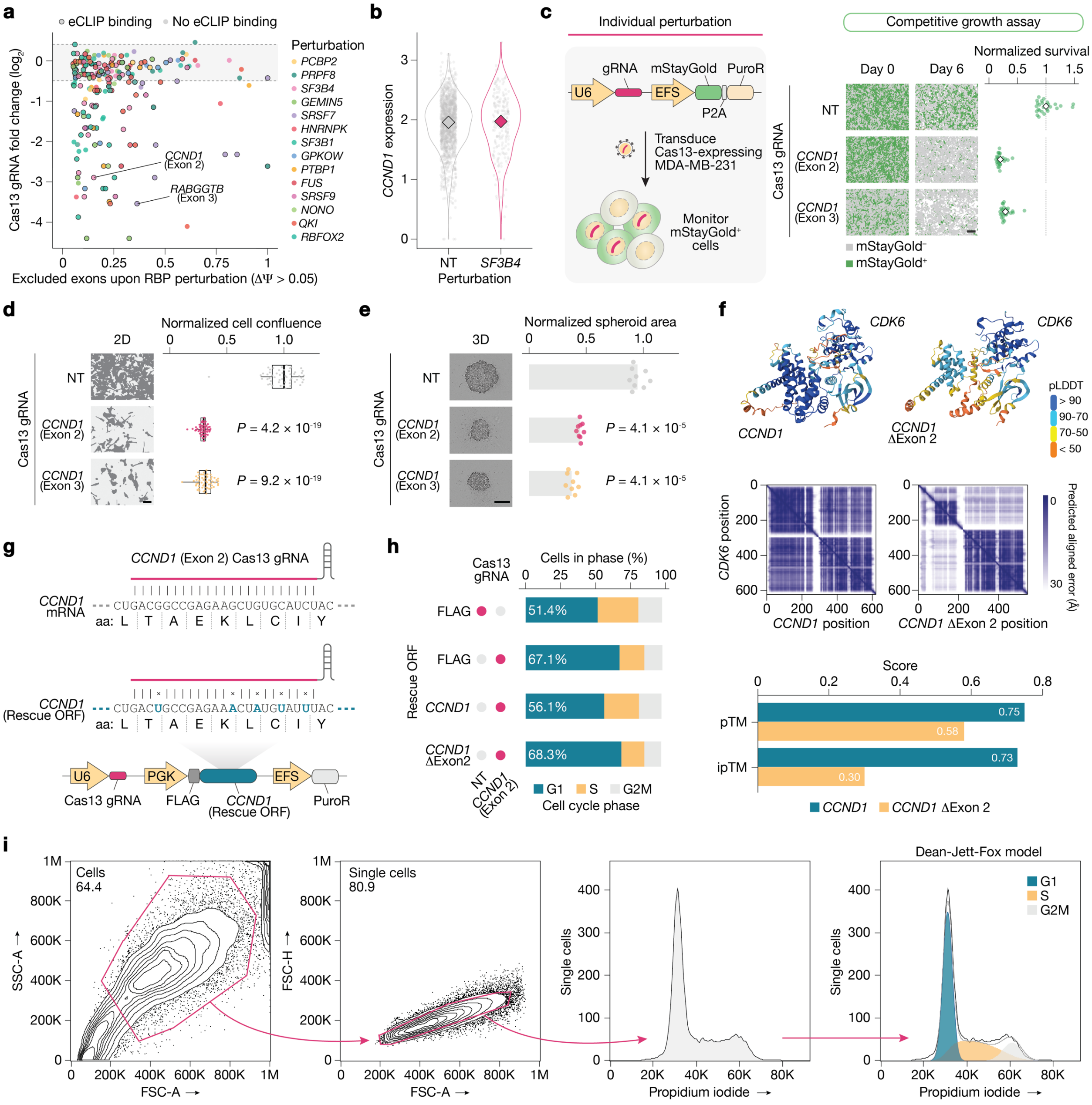


**Supplementary Fig. 4: Exon skipping in *CCND1* leads to a loss-of-function isoform.**
**a,** *Rfx*Cas13d guide RNA (gRNA) fold change (day 14 vs. day 0) from Liang et al.[5] targeting exons of transcripts that are excluded upon RBP perturbations in CRISPore-seq. **b,** Single-cell gene expression of *CCND1* in *SF3B4*-perturbed cells or cells that received non-targeting (NT) control perturbations. **c,** Competitive growth assay to quantify the impact of individual Cas13 perturbations on cell fitness. Representative images and normalized survival (day 6 vs. day 0) of mStayGold-positive *Rfx*Cas13d MDA-MB-231 cells transduced with gRNAs targeting *CCND1* exon 2, exon 3, or a non-targeting gRNA ($n$ = 27 images from three independent transductions). **d,** Representative images and normalized cell confluence of *Rfx*Cas13d MDA-MB-231 cells transduced with gRNAs targeting *CCND1* exon 2, exon 3, or a non-targeting gRNA at 4 days after doxycycline-induction ($n$ = 54 images per timepoint with three images per technical replicate, six technical replicates per transduction, and three independent transductions per perturbation).

Boxplots show median, interquartile range, and whiskers indicating 1.5× interquartile range. Significance was determined using a two-sided Mann-Whitney *U*-test. Scale bar, 100 µm. **e,** Representative images and normalized spheroid area of *Rfx*Cas13d MDA-MB-231 cells transduced with gRNAs targeting *CCND1* exon 2, exon 3, or a non-targeting gRNA at 7 days after doxycycline-induction ($n = 9$ images with one image per technical replicate, three technical replicates per transduction, and three independent transductions per perturbation). Significance was determined using a two-sided Mann-Whitney *U*-test. Scale bar, 400 µm. **f,** Structural predictions (*top*) for the CCND1-CDK6 or CCND1(ΔExon 2)-CDK6 complex formation using AlphaFold3[6] with the per-residue confidence score using the predicted Local Distance Difference Test (pLDDT). Heatmaps show the predicted aligned error (*middle*). Template modeling (pTM) and interchain predicted template modeling (iPTM) scores for the CCND1-CDK6 or CCND1(ΔExon 2)-CDK6 complex models (*bottom*). **g,** Silent mutations in the second exon of *CCND1* for phenotypic rescue assays to escape Cas13 knockdown. **h,** Cell cycle phase distribution of MDA-MB-231 cells after perturbation of endogenous *CCND1* and/or rescue with FLAG, FLAG-*CCND1*, or FLAG-*CCND1*(ΔExon 2) expression at 4 days after doxycycline-induction. The bars denote the mean of three independent transductions. **i,** Gating strategy and fitting for cell cycle assays of MDA-MB-231 cells. Cells were gated for singlets and the cell cycle phase distribution was computed using Dean-Jett Fox model fits on the propidium iodide signal intensity.

### Supplementary References

1. Barry, T., Mason, K., Roeder, K. & Katsevich, E. Robust differential expression testing for single-cell CRISPR screens at low multiplicity of infection. *Genome Biol* **25**, 124 (2024).
2. Tsherniak, A. *et al.* Defining a Cancer Dependency Map. *Cell* **170**, 564-576.e16 (2017).
3. Ashburner, M. *et al.* Gene Ontology: tool for the unification of biology. *Nat Genet* **25**, 25–29 (2000).
4. Milacic, M. *et al.* The Reactome Pathway Knowledgebase 2024. *Nucleic Acids Res* **52**, D672–D678 (2024).
5. Liang, W.-W. *et al.* Transcriptome-scale RNA-targeting CRISPR screens reveal essential lncRNAs in human cells. *Figshare preprint* (2025)
6. Abramson, J. *et al.* Accurate structure prediction of biomolecular interactions with AlphaFold 3. *Nature* **630**, 493–500 (2024).
